# Empowering systems-guided drug target discovery with metabolic and structural analysis

**DOI:** 10.1101/2021.05.17.444532

**Authors:** Sourav Chowdhury, Daniel C. Zielinski, Christopher Dalldorf, Joao V Rodrigues, Bernhard O. Palsson, Eugene I Shakhnovich

**Author notes:** Contributed Equally (Sourav Chowdhury, Daniel C. Zielinski).

## Abstract

Elucidating intracellular drug targets has been a difficult problem. While machine learning analysis of omics data has been a promising approach, going from large-scale trends to specific targets remains a challenge. Here, we developed a systems-guided hierarchic workflow that utilizes metabolic and structural analysis to narrow in on specific targets suggested by statistical and machine learning analysis of metabolomics data. Utilizing a novel multi-valent DHFR-targeting antibiotic compound, CD15-3, as a case study, we first measured global metabolomics and applied statistics and machine learning to locate broad areas of metabolic perturbation under antibiotic stress. We then tested the ability of suggested compounds to rescue growth and performed metabolic modelling to identify pathways whose inhibition was consistent with growth rescue patterns. Next, we utilized protein structural similarity to further prioritize candidate drug targets within these pathways. Overexpression and *in vitro* activity assays of a top candidate target, HPPK (folK), showed complete recovery from drug induced growth inhibition and with microscopy. As interest in ‘white-box’ machine learning methods continues to grow, this study demonstrates how established machine learning methods can be combined with mechanistic analyses to improve the resolution of drug target finding workflows.

## Introduction

Pharmaceuticals are often used with incomplete knowledge of their intracellular drug binding partners (Silver, 2007). For example, while antibiotic molecules are designed to selectively inhibit essential bacterial proteins (Butler et al., 2013), even conventional antibiotic drugs are found to target multiple molecular targets inside bacterial cells. Identifying the full spectrum of drug targets is critical to understanding drug mechanism of action as well as to exploit multivalency to tackle the problem of drug resistance.

Recent advances in systems biology tools are making systematic search for intracellular drug targets an increasingly accessible task (Rabinowitz et al., 2011). Large-scale measurement of drug perturbation, such as metabolomics or transcriptomics, coupled to machine learning has been a promising approach to understanding drug mechanism of action and targeting. These methods can identify trends that are uniquely associated with a drug effect. However, identifying specific targets from machine learning is challenging due to the difficulty in interpreting such models. By contrast, mechanistic models and targeted analyses have the advantage of greater interpretability but struggle to learn from large-scale datasets.

In this work, we developed a systems-guided multiscale drug target finding workflow that integrates machine learning analysis of metabolomics data with metabolic modelling and protein structural analysis. First, we analyse untargeted global metabolomics with statistics and machine learning to unravel differences in the global metabolome upon antibiotic treatment and captured potential hotspots of metabolic perturbation. We then integrate metabolic supplementation growth rescue experiments with metabolic modelling to identify metabolic pathways whose inhibition is consistent with data. Finally, we perform protein structural analysis to identify likely targets within candidate pathways based on similarity to known targets and validate these candidates experimentally.

We deployed this workflow to identify the functional target of CD15-3, a novel antibiotic compound previously developed in house (Zhang et al., 2021). CD15-3 was designed to interact with wild type DHFR (dihydrofolate reductase) and its trimethoprim (TMP) resistant mutants (Zhang et al., 2021). In our previous report we showed that CD15-3 interacts with DHFR; however, overexpression of the DHFR-encoding gene folA was only able to partially rescue CD15-3-induced growth inhibition. The lack of complete growth rescue from overexpression of DHFR indicated the presence of an additional non-DHFR intracellular target of CD15-3, which could be responsible for the growth inhibitory effect of the compound (Rodrigues and Shakhnovich, 2019a; Zhang et al., 2021).

Here, we generate metabolomics data for cells treated with CD15-3 and analysed the data with a combination of machine learning, metabolic modelling, and structural analysis to suggest priority candidates for alternate binding targets. We then experimentally validate the inhibition of top candidate targets by CD15-3 using gene overexpression, imaging experiments and in vitro enzyme assays. We propose that this systems-guided multi-scale framework integrating multiple interconnected layers to identify the intracellular target can be potentially used to understand intracellular mode of action and molecular targets of other novel antibiotic formulations as well as other drug candidates with unknown target.

## Results

### An integrated metabolomics-guided framework to identify intracellular drug targets

Antibiotic molecules often have unintended intracellular target. A systems wide analysis capturing the drug induced intra-cellular perturbation can be insightful in unravelling the intracellular mechanism of action of the drug and its molecular target. We developed an untargeted global metabolomics-guided multi-layered analysis framework for antibiotic target identification (Figure 1).

**Figure 1:**
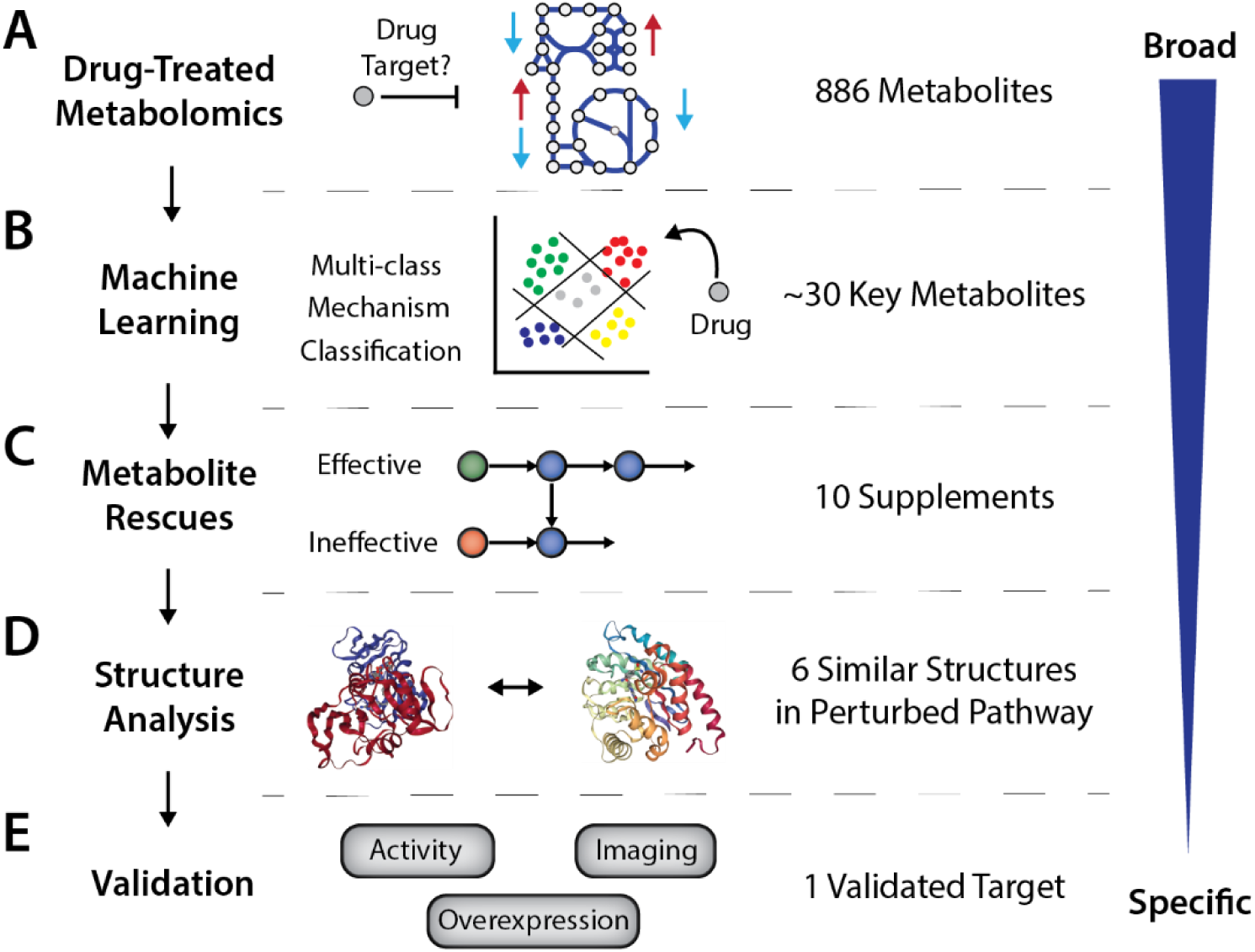
Metabolomics-guided drug target finding workflow. **(A)** Cells are treated with the compound of interest, and metabolite changes are measured with high throughput metabolomics. **(B)** Machine learning is used to identify drug-specific perturbations by comparison to publicly available drug response metabolomic profiles. **(C)** Key perturbed metabolites are provided as supplements to attempt growth rescue, and metabolic modeling is used to analyze the results to determine whether patterns are consistent with inhibition of certain metabolic pathways by the compound. **(D)** Structural analysis identifies candidate enzymes through homology to known targets of the drug. **(E)** Candidate genes are validated through a combination of overexpression of the target to rescue growth, activity assays to verify binding and inhibition of the target, and imaging to determine phenotypic effect.

This workflow analyzes drug-treated metabolomics data using a combination of machine learning, metabolic modeling, and protein structures to prioritize candidate targets of antimicrobial inhibition. First, we perform an untargeted metabolomic analysis to identify metabolites that are highly perturbed by the drug, obtaining a broad assessment of drug activity, as well as evaluating the ability of these key metabolites to rescue growth (Figure 1A). Independently, we utilize machine learning and a previously published dataset of metabolomic response of diverse antibiotics(Zampieri et al., 2017) to identify mechanism-specific and unique drug signatures in the metabolomic response (Figure 1B). Then, we utilize metabolic modeling to identify metabolic pathways that when inhibited would result in measured patterns of growth rescue (Figure 1C). Finally, we perform a structural analysis of global and active site properties to the intended target of the drug to identify likely off targets (Figure 1D). Based on the evidence of these three analytical tools, we prioritized candidate targets for experimental validation using gene overexpression, enzyme assays, and cell imaging to discern phenotypic changes (Figure 1E).

We deployed this framework as a case study to search for the intracellular mode of action of a novel evolution drug lead CD15-3 (Zhang et al., 2021) and determine its non-DHFR target.

### Metabolomic analysis of CD15-3 perturbation

We first measured the metabolic perturbation upon CD15-3 treatment to obtain a metabolome-wide view of CD15-3 action inside the cell. To that end we carried out untargeted global metabolomics measurements to obtain the comparative global metabolome under untreated and CD15-3 treated conditions (Figure 2). Cells were grown in the presence and absence of CD15-3 for different lengths of time and were harvested for processing. Comparative metabolite abundances showed progressively increasing differences in cellular metabolism as the cells were exposed to CD15-3 for longer lengths of time. Figure 2A shows the differences in metabolite abundances involved in nucleotide metabolism, carbohydrate metabolism, cofactors, and peptides. As observed, a five-hour exposure to CD15-3 significantly impacts the global metabolism of cells, with even greater differences being observed after 12 hours exposure.

**Figure 2:**
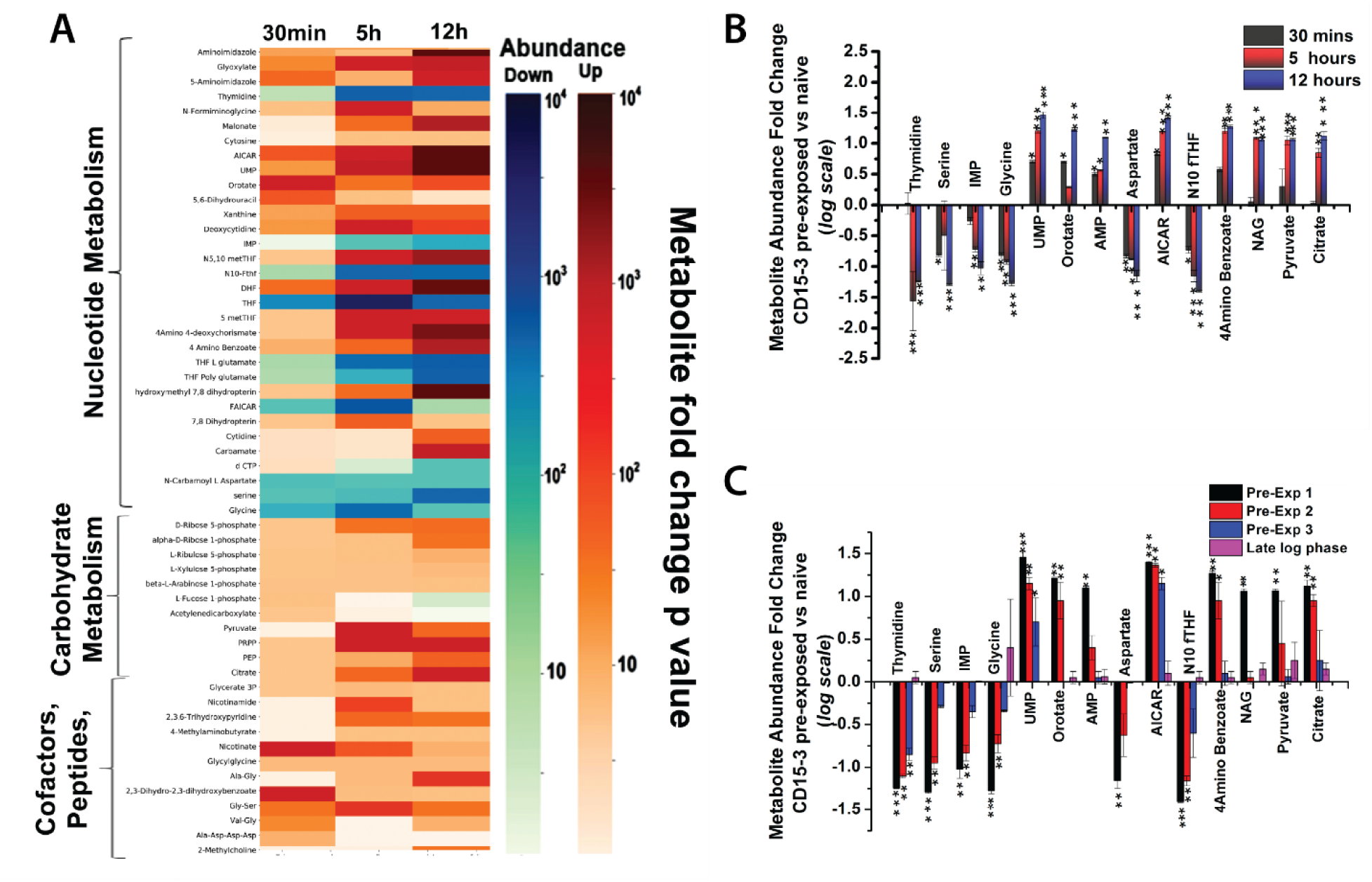
Metabolomics data following perturbation with the antibiotic candidate CD15-3. **(A)** Heat map describing the global metabolomic profile of untreated (WT) and treated (WT+CD15-3) cells. Color gradient indicates fold differences (in log scale) observed in the metabolite abundance levels in the CD15-3 treated set compared to the untreated WT cells. Horizontal lines in the heat map indicate metabolites. Color bar shows negative logarithm of p-values (Mann-Whitney Test) associated with abundance ratios (treated vs. control) of each metabolite. A higher intensity of the blue bar indicates more depletion while a higher intensity of the orange bar indicates more accumulation. **(B)** Bar plot showing abundance fold differences (log scale) of selected metabolites as the cells grow in the presence of CD15-3. Metabolomics data for the treated and untreated control cells were gathered by harvesting cells at three different time points as shown in the figure. **(C)** Bar plot showing the metabolite abundance profile (log scale) of selected metabolites under the condition of the recovery experiment when cells pre-incubated with CD15-3 were grown in normal M9 media supplemented with 0.8g/L glucose. The fold differences in the metabolite abundances are the ratios of metabolite abundance observed in the (recovering) pre-exposed/CD15-3 treated cells to that of the naïve cells. (* indicates metabolite abundance ratios with p values ≤0.05, ** ≤0.001, *** ≤ 0.0001 as derived from a Mann-Whitney test).

Thymidine, a constituent of the pyrimidine biosynthesis pathway shows a 15-fold drop in abundance after 5 hours of growth with CD15-3 treatment and a 17-fold drop after 12 hours of growth (Figure 2B). 4-aminobenzoate which is synthesized from chorismate and is an important metabolic intermediate leading to the biosynthesis of a host of crucial metabolites such as folates, shows 15-fold higher abundance at 5 hours of growth and 18-fold higher abundance at 12 hours of growth with CD15-3 treatment. N10-formyl-THF, a precursor in the THF biosynthesis, showed 12-fold up regulation at 5 hours of growth with CD15-3 treatment and 15-fold higher abundance after 12 hours growth. Folates are crucial for the biosynthesis of many important cellular metabolites, including glycine, methionine, formyl-methionine, thymidylate, pantothenate and purine nucleotides. Our comparative global metabolome showed significant fold differences in many of these metabolites indicating some plausible perturbation around the folate pathway and gross perturbation distributed throughout the overall nucleotide metabolism. For example, serine and glycine showed continuous cellular depletion upon CD15-3 treatment with more than 20-fold lower abundances after 12 hours of growth under CD15-3 treatment. Another interesting metabolic marker for perturbed purine metabolism is AICAR, which showed almost 8-fold higher abundance at 30 minutes of growth and 16-fold higher abundance at 5 hours of growth. Cellular buildup of AICAR at early stages of treatment could indicate that purine metabolism gets disrupted quite early under CD15-3 treatment. UMP, a constituent of pyrimidine metabolism also showed cellular build up with 32-fold higher abundance after 12 hours of growth with treatment. Also, significant fold differences in the abundance levels of various peptides, cofactors and lipids were observed which too could be attributed to a CD15-3 induced metabolic stress response (Zhang et al., 2021). We observed significant fold differences in some metabolites constituting carbohydrate metabolism. Pyruvate and citrate cellular buildup has been known to be associated with metabolic stress response (Shimizu, 2014). Under CD15-3 treatment we observed 11-fold higher abundance of pyruvate and 12-fold higher abundance of citrate at 12 hours growth with CD15-3 treatment. Cellular buildup of citrate under CD15-3 treatment potentially indicates a possible slowdown of glycolysis and in turn energy metabolism.

We went on to determine the abundance levels of the relevant metabolites during recovery following CD15-3 treatment. To determine which metabolites display the most delayed recovery after CD15-3 treatment and hence are most impacted upon the treatment, we incubated WT cells in CD15-3 for 12 hours and subsequently transferred them to M9 media supplemented with 0.8g/L glucose. In a parallel control set, WT cells were grown for 12 hours in M9 media and subsequently regrown in the same media. Metabolite abundance levels were measured at four discrete time points (Supplementary Figure 1) of cell harvesting. Notably, AICAR and several other metabolites that showed significant abundance changes in early stages of treatment gradually restored to normal levels during recovery (Figure 2C). However, thymidine, IMP, and serine had significant fold differences until the last pre-exponential phase (which we term pre-exponential phase 3 as shown in Supplementary Figure 1 as PE3), which occurred between 3 and 5 hours. A similar trend was observed with AICAR, UMP, and N10-formyl-THF, as these metabolites took longer to recover. N-acetylglutamate (NAG), which is a constituent of the ornithine biosynthesis pathway via the formation of N-acetyl ornithine, had significantly higher abundance upon treatment and was found to respond much earlier in the recovery experiment with abundance levels quickly returning to normal.

### Machine learning reveals antibiotic mechanism-specific perturbations

To aid the interpretation of the measured metabolomics data, we developed a machine learning workflow to identify metabolic signatures associated with both known and unknown targets of a compound. To contextualize the metabolomic response for CD15-3 and separate drug-specific effects from general growth inhibitory effects, we utilized a previously published survey of the metabolomic response of *E.* coli to diverse antibiotics (Zampieri et al., 2017). We trained a multi-class logistic regression (LR) model to identify metabolomic perturbations associated with each of five possible mechanisms (antifolate, cell membrane, DNA synthesis, translation, oxidative stress) (Figure 3A). Visualizing the data with Uniform Manifold Approximation and Projection (UMAP) and clustering the projection revealed that CD15-3 perturbation showed similarity to several other antibiotics including the DHFR-targeting antibiotic trimethoprim, as well as hydrogen peroxide perturbation that was used as a control to approximate a generic antibiotic growth inhibition (Figure 3B). The multi-class LR model performed well for antifolates (Figure 3C), the class of greatest interest, although it performed poorly for many other antibiotic classes. Consistent with the UMAP projection, the LR model suggests that while CD15-3 shows characteristics as an antifolate at early time points, it shows a broader growth inhibitory response consistent with non-specific antibiotic perturbation (Figure 3D).

**Figure 3.**
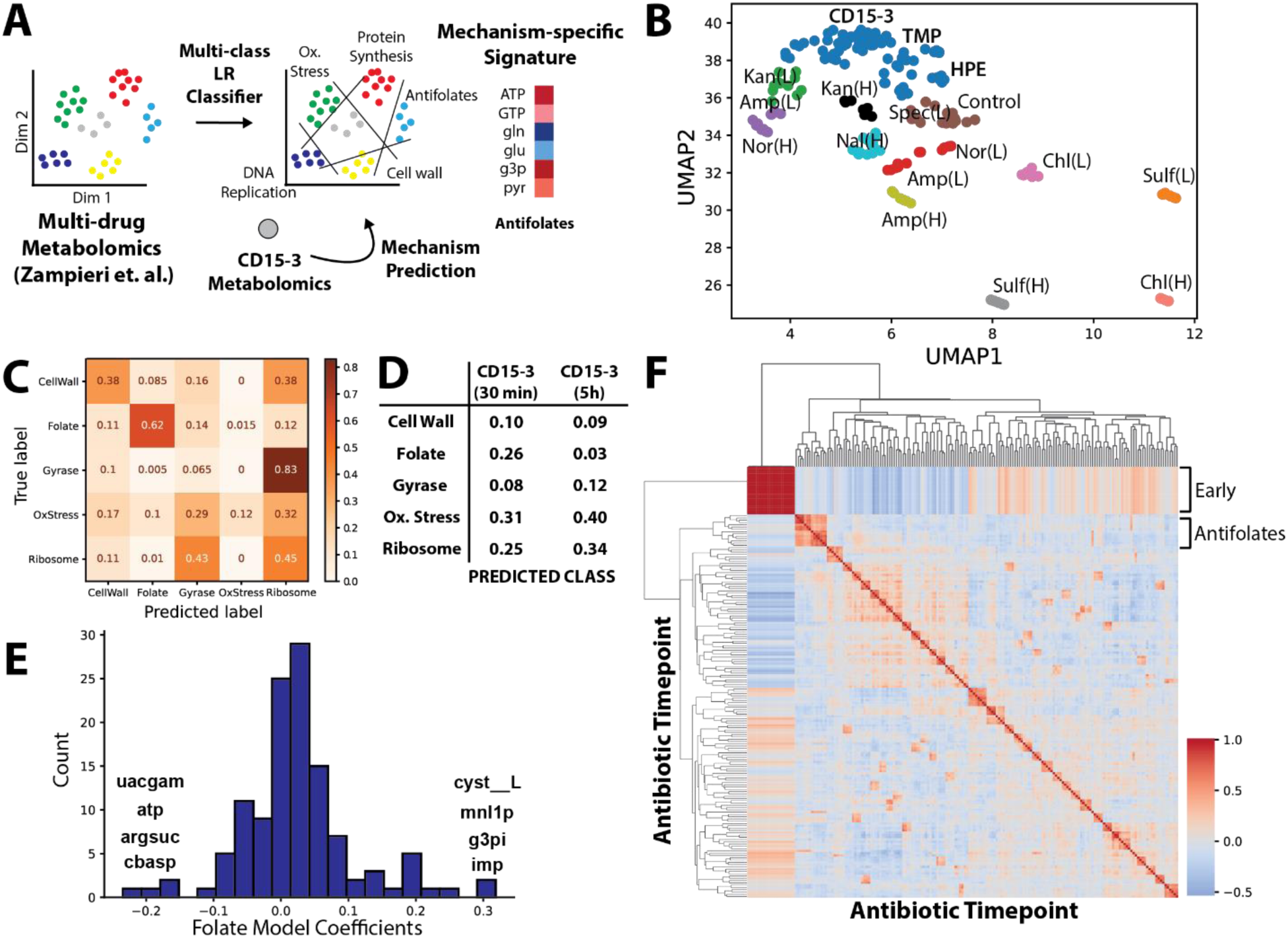
Machine learning identifies mechanism-specific trends in metabolomics data. A) A multi-class Logistic Regression model was used to identify signatures associated with 5 different drug classes from a previous study (Zampieri et. al. 2017) This signature is removed from the CD15-3 metabolomics data to calculate a residual drug-specific signature that is examined for signs of additional targets. B) Uniform Manifold Approximation and Projection (UMAP) plot of metabolomics data with clusters (N=21) generated by Density-based spatial clustering of applications with noise (DBSCAN). C) Confusion matrix demonstrates multi-class prediction accuracy for the antifolates class of antibiotics. D) Predicted drug mechanism from the logistic regression model for CD15-3 at 30-minute and 5-hour exposure. E) Model feature importance shows diverse metabolites contributing to the antifolate signature. F) Correlation matrix of filtered metabolomics data shows that most drugs induce highly disparate perturbations as evidenced by lack of clustering behavior of samples, with exception to the antifolate class.

We further evaluated this model to identify metabolites that were key to distinguishing the antifolate response (Figure 3E). While metabolite scores contributing to the antifolate class prediction were diverse, top hits included several purine and pyrimidine adjacent metabolites (ATP, IMP, Argininosuccinate, Carbamoyl-L-aspartate) consistent with antifolate inhibition of nucleotide metabolism. Among these, ATP and IMP were identified as perturbed from the statistical analysis of CD15-3 response, and these were utilized as targeted for growth supplementation experiments in the next section.

Investigating markedly better performance of the antifolate model over other classes of antibiotics, we found that correlation of metabolomic response is substantially greater across different antifolates compared with other classes of drugs (Figure 3F). The consistent perturbation seen in antifolates is logical given the critical metabolic role of the folate pathway in growth.

### Metabolic modeling predicts patterns in growth rescue experiments for candidate pathway inhibitions

The metabolite response to CD15-3 identified major perturbations in the nucleotide metabolism. Further the statistical and machine learning analysis revealed that CD15-3 metabolomic perturbation has both antifolate and generic antibiotic response signatures. To further narrow down which perturbations are most directly linked to CD15-3-dependent growth inhibition, we chose to use a subset of the identified metabolic markers for metabolic supplementation experiments to attempt a rescue from CD15-3-induced growth inhibition (Figure 4A). To this end, wild type cells were grown in the presence of the externally supplemented metabolites under conditions of CD15-3 treatment. These externally supplemented metabolites were selected owing to their highly perturbed abundance levels in the comparative global metabolome (CD15-3 treated versus untreated) and their proximity to folate pathway, as DHFR was the intended target for CD15-3 (Zhang et al., 2021).

**Figure 4:**
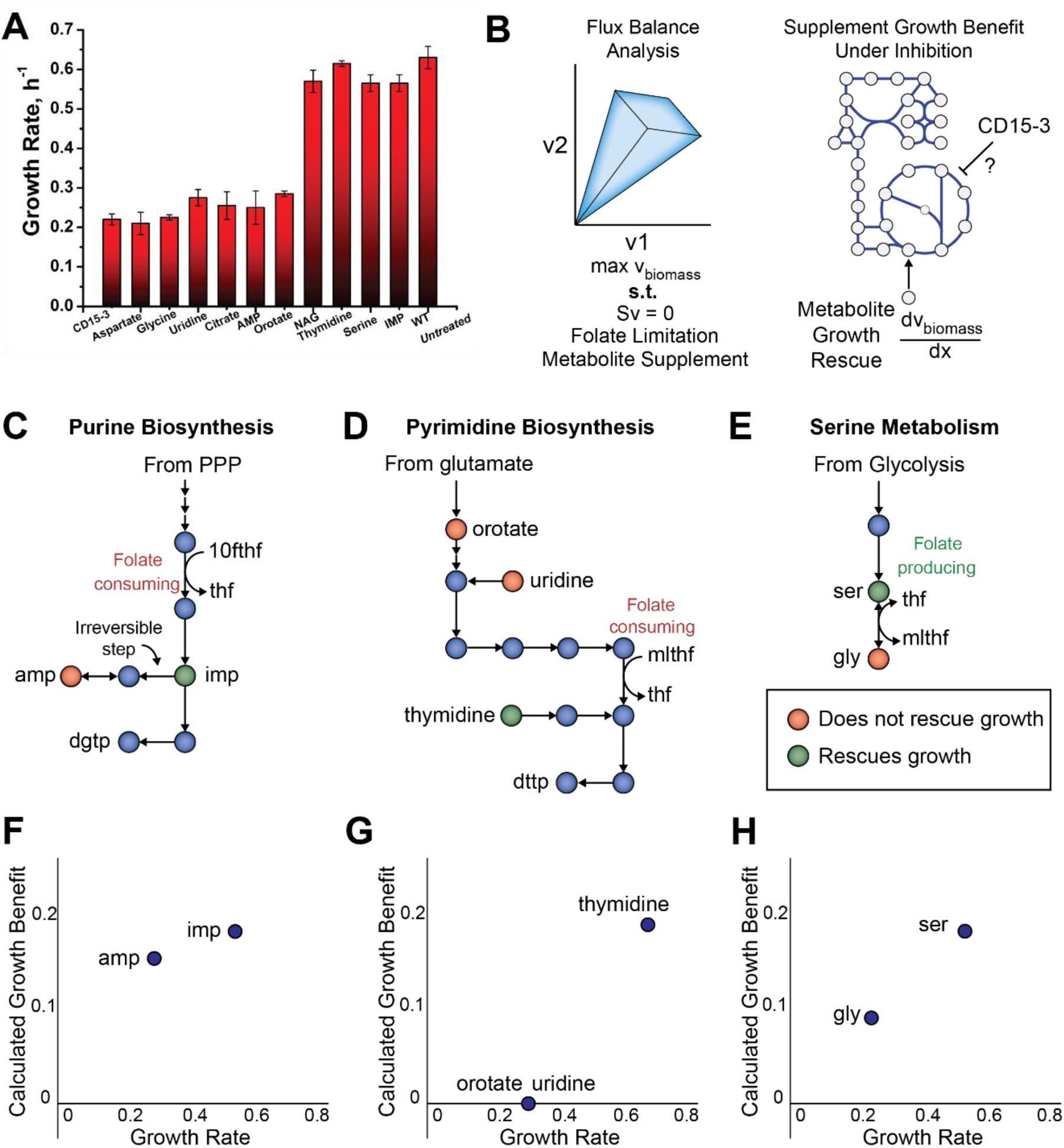
Effect of nutrient supplements on CD15-3-treated cells indicate broad inhibition of folate pathway. **(A)** Bar plot showing how metabolite supplementation impacts growth rate in the presence of CD15-3. Rescue from CD15-3-induced growth inhibition is observed with thymidine, serine, NAG and IMP supplementations. A fixed concentration of 200µM CD15-3 was used in these supplementation experiments. **(B)** A metabolic supplementation strategy was employed for target identification by utilizing metabolic pathway analysis to find supplementation patterns consistent with inhibition of a particular metabolic pathway. Flux balance analysis was used to identify metabolic pathways with inhibition consistent with observed growth rescue patterns. **(C**) Purine biosynthesis pathway. Both IMP and AMP supplements are downstream of folate-requiring steps. However, AMP is unable to be converted to dGTP due to an irreversible step in the pathway, while IMP can be converted to both dATP and dGTP. **(D)** Pyrimidine biosynthesis pathway. The growth rescuing supplement, thymidine, enters the pathway downstream of the folate requiring step. Supplements that enter upstream of the folate requiring step, namely orotate and uridine, did not rescue growth. **(E)** Serine and glycine biosynthesis. Serine, which produces the charged folate form 5,10-methylene tetrahydrofolate when converted to glycine, rescued growth, while glycine itself did not rescue growth. **(F)** Correlation of growth rate from different supplements with purine biosynthesis metabolites and corresponding model calculated growth benefit**. (G)** Correlation of growth rate from different supplements with pyrimidine biosynthesis metabolites and corresponding model calculated growth benefit**. (H)** Correlation of growth rate from different supplements with serine metabolism metabolites and corresponding model calculated growth benefit.

In all these experiments, the external metabolite supplement concentration was kept at 0.5mM. Control experiments were performed in the absence of CD15-3 to check for intrinsic toxicity of these external metabolite supplementations (Supplementary Figure 2A). We observed that the metabolites selected for the metabolic supplementation experiments did not show toxic effect on the bacterial growth. Also, we checked the possibility that these compounds might serve as alternate carbon sources. To that end we grew cells without CD15-3 in M9 minimal media with each of these supplements (without glucose). The growth profile was compared with bacterial growth in M9 minimal media with 0.8g/L glucose, which is the typical media composition used in the rest of the study. We observed that, apart from thymidine, no other metabolite served as an alternate carbon source (Supplementary Figure 2B). In the presence of thymidine, cells did show slow growth with a very long lag phase compared with the glucose control.

Supplementation with thymidine, NAG, serine, and IMP showed growth rescue, as reflected in improved growth rates (Figure 4B) under the conditions of CD15-3 treatment. On the other hand, negligible or no effect on the growth rates were observed with external supplementation of aspartate, glycine, uridine, citrate, orotate and AMP. Thymidine and other metabolites which showed significant rescue of CD15-3 inhibition improved the growth rate to around 0.6 h^-1^ which is comparable to the growth rates observed in the WT cells in the absence of CD15-3.

It is interesting to note that metabolites whose external supplementation rescued growth rates did not significantly affect the lag time (Supplementary Figure 2C). Lag time in bacterial growth is a critical indicator of cellular adaptation in the ambient growth condition (Fridman et al., 2014). In our experimental condition a higher lag time in the presence of some external supplements and CD15-3 treatment reflects time of metabolic rewiring and adaptation in the supplement enriched growing condition. It is interesting to note that NAG and serine significantly prolonged the lag time (Supplementary Figure 2C) under treatment conditions although both have shown to have positive impact in improving growth rates (Figure 4B). Thus, NAG and serine could be considered as partial rescuers from CD15-3 induced stress with improvement on only growth rate and worsening the lag time.

Metabolic supplementation demonstrated that diverse compounds were able to rescue inhibition by CD15-3, while other metabolites had no effect. To better understand the metabolic rationale for the supplementation rescue patterns of CD15-3, we utilized the most updated metabolic network reconstruction of *E. coli*, iML1515 (Monk et al., 2017). We examined the trends in metabolite supplementation rescue experiments with respect to their location in corresponding metabolic pathways. We observed that the effectiveness of the supplement in rescuing growth was determined by the position of the supplement in the metabolic network viz a viz folate metabolism. Specifically, supplements that have the potential to mitigate folate deficiency, namely thymidine, IMP, and serine, were effective at rescuing growth. Supplementation with Thymidine and IMP bypass the folate-dependent step in pyrimidine and purine biosynthesis, respectively (Figure 4C, D), while serine can contribute to folate production through its conversion to glycine (Figure 4E). Meanwhile, supplements that do not mitigate a folate deficiency do not rescue growth, even though these metabolites may be adjacent in the network to successful supplements, such as uridine, AMP, and glycine. Uridine is a pyrimidine precursor that is upstream of the folate-dependent biosynthetic step. Glycine has an unclear role, as metabolism through the glycine cleavage chain produces folate, while conversion to serine consumes folate. Thus, although DHFR may not be the primary target of CD15-3, the inhibitory activity of CD15-3 still appears to primarily work through folate limitation.

To rigorously evaluate the hypothetical effect of these supplements, we developed a constraint-based metabolic modelling workflow to assess whether inhibition of a particular pathway is consistent with observed growth rescue patterns of supplements (Figure 4B, Methods). Constraint-based modelling through flux balance analysis utilizes a metabolic network reconstruction to predict reaction flux through the metabolic network that maximizes growth for a given experimental condition (Orth et al., 2010). We evaluated two possible metabolic inhibition scenarios: direct pathway inhibition, and cofactor depletion (see Methods). To evaluate possible direct pathway inhibition, we computationally inhibited metabolic reactions one by one and calculated the ability of each metabolite in the model to rescue growth. To evaluate possible cofactor depletion, we generated cofactor draining reactions and calculated the ability of each metabolite in the model to generate additional cofactor charges. Finally, we correlated the model-calculated benefit of each metabolite to the experimental observed growth rescue potential of those metabolites.

We found that the experimental growth rescue pattern was most consistent with a folate cofactor drain mechanism of CD15-3 (Figure 3G). Comparing the model-calculated to the observed growth benefit of different metabolites revealed an ability of the model to distinguish the benefit of similar metabolites. For example, the improved ability of IMP over AMP to rescue growth (Figure 4F), the improved ability of thymidine over uridine to rescue growth (Figure 4G), and the ability of serine but not glycine to rescue growth (Figure 4H), were all predicted correctly by the model, after accounting for wild type enzyme expression as detailed below.

We note that the metabolic model did not inherently incorporate the wild type expression state under which supplements were administered. To correct for wild type gene expression, we shut off flux through several reactions based on measured lack of expression under wild type conditions (Schmidt et al., 2016). For example, the model initially predicted citrate to have a growth rescuing effect; however, the citrate transporter is not expressed under normal conditions. Similarly, the model calculated only a minor benefit of IMP over AMP initially, but investigation revealed that the AMP incorporation pathway utilized by the model included both spontaneous reactions, which are not likely to occur at a high enough rate to sustain growth, and the cryptic gene adeD, encoding adenine deaminase. These cases were handled individually utilizing available expression data (Schmidt et al., 2016), and all model corrections are included in the metabolic modelling code in the Supplementary Materials.

To determine whether the observed agreement was specific to folate inhibition or was associated with growth more broadly, we additionally implemented random pathway inhibitions and compared the growth benefit under these conditions. While some metabolites, such as serine and glycine, had growth rescue behaviour that agreed more broadly with general growth benefit, other metabolites such as AMP/IMP and thymidine/uridine agreed substantially better with folate inhibition than inhibition of other pathways (Supplementary Figure 3). Thus, the metabolic modelling results were consistent with topological pathway analysis to point to folate inhibition as the key metabolic limitation induced by CD15-3, despite DHFR being ruled out as the sole growth limiting enzyme by previous work (Zhang et al., 2021).

### Structural analysis of possible alternate binding targets

Experimental evidence from metabolomics supplementation, all suggested that folate perturbation is the primary mode of action of CD15-3. Thus, we hypothesized that the alternate target of CD15-3 also lies within the same metabolic pathway as DHFR. Notably, other enzymes in the folate biosynthetic pathway are also known to be drug targets, such as Dihydropteroate synthase, the target of sulfamethoxazole. To narrow down likely alternative targets of CD15-3, we employed a genome-scale reconstruction of enzyme protein structures, termed the GEM-PRO or Genome-scale Model with Protein Structures of *E. coli* (Monk et al., 2017) (Figure 5A). We first computed the overall similarity of global structural properties of all protein chains in the GEM-PRO, which showed DHFR and other folate pathway enzymes clustered separately from the majority of metabolic enzymes (Figure 5B). We found that DHFR did not have a high degree of structural similarity to any particular enzyme based on whole chain property similarity (Figure 5C). Utilizing the FATCAT algorithm for structural alignment (Li et al., 2020a) and isomif analysis of annotated binding sites, we compared aligned structural similarity of the intended target DHFR and other enzymes in the folate biosynthetic pathway (Figure 5D). While similarity of whole chains was generally low (Figure 5E), comparison of active sites suggested possible alternative binding targets for CD15-3 (Figure 5F). Notable proteins with high binding pocket similarity scores included several enzymes involved in folate biosynthesis and folate interconversion, such as MTHFC, HPPK2, and DHPS. Thus, we decided to screen several candidate upstream enzymes in the folate biosynthetic pathway whose binding pockets have high similarity to DHFR.

**Figure 5.**
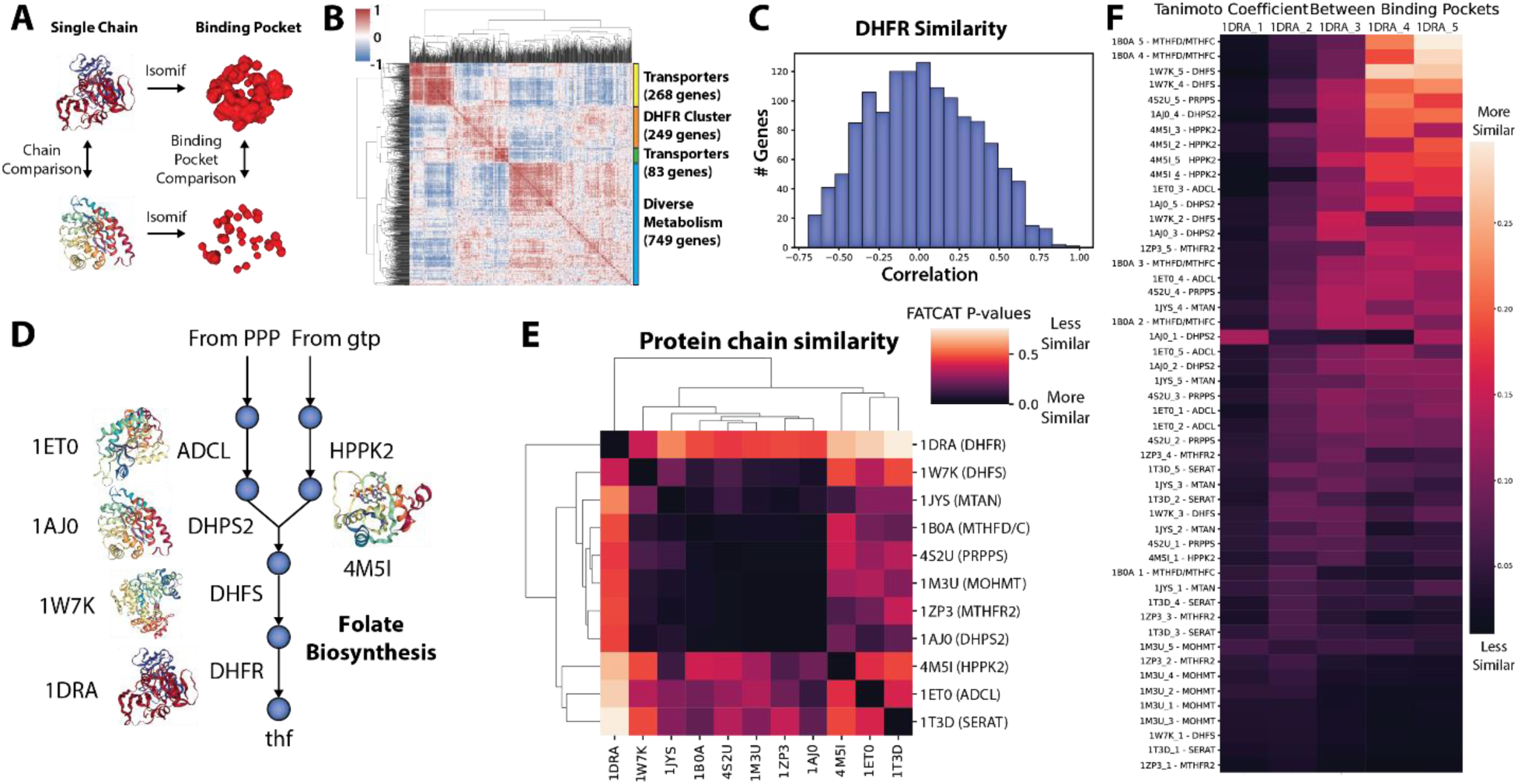
Protein structural similarity evidence supports other folate enzymes as possible targets of CD15-3. **(A)** Overview of workflow for structural analysis. **(B)** Global pairwise structural similarity (Pearson correlation R) across metabolic proteins based on calculated structural properties from the E. coli GEM-PRO. **(C)** Global structural similarity (Pearson correlation R) of DHFR, the intended target of CD15-3, to other genes. **(D)** Overview of folate pathway and protein structures extracted from PDB. **(E)** Results for whole structure similarity analysis utilizing FATCAT. Low p-values reported by FATCAT indicate more similar structures. **(F)** Results for active site analysis using isomif for comparison. Protein structure identifiers are taken from PDB.

### In vivo validation of the intracellular target of CD15-3

Utilizing the prioritized target list from structural analysis, we evaluated whether overexpression of any of these enzymes rescued growth inhibition by CD15-3. Regulated overexpression of candidate proteins viz., HPPK (encoded by gene folK), DHPS (encoded by gene folP), DHFS (encoded by folC), MTHFC (encoded by gene folD), MTHFR (encoded by gene metF) and ADCL (encoded by gene pabC) using the pBAD promoter with 0.1% arabinose induction was carried out to determine whether any of the overexpressed genes show recovery from CD15-3 induced growth inhibition and thus may be the non-DHFR target of CD15-3. In our previous study (Zhang et al., 2021) we showed that overexpression of folA (encoding DHFR) partially rescued CD15-3 induced toxicity at lower concentrations of CD15-3.

Of the assessed proteins only overexpression of folK showed clear sign of rescue from growth inhibition (Figure. 6A). folK encodes for 6-Hydroxymethyl-7,8-dihydropterin pyrophosphokinase (HPPK). HPPK is a key enzyme in the folate biosynthesis pathway catalyzing the pyrophosphoryl transfer from ATP to 6-hydroxymethyl-7,8-dihydropterin, is an attractive target for developing novel antimicrobial agents (Bermingham and Derrick, 2002; Chhabra et al., 2013; Chhabra et al., 2012). Upon folK overexpression cells did not show any change in growth rates in a broad range of concentrations of CD15-3 (Figure 6A). It stays close to 0.6 h^-1^ at all the concentrations of CD15-3, which is the typical growth rate of WT in the absence of CD15-3. On the other hand, overexpression of folA at 0.005% arabinose induction showed only partial rescue from CD15-3 inhibition (Figure 6B). Rescue in growth rate was more pronounced at lower CD15-3 concentration.

**Figure 6:**
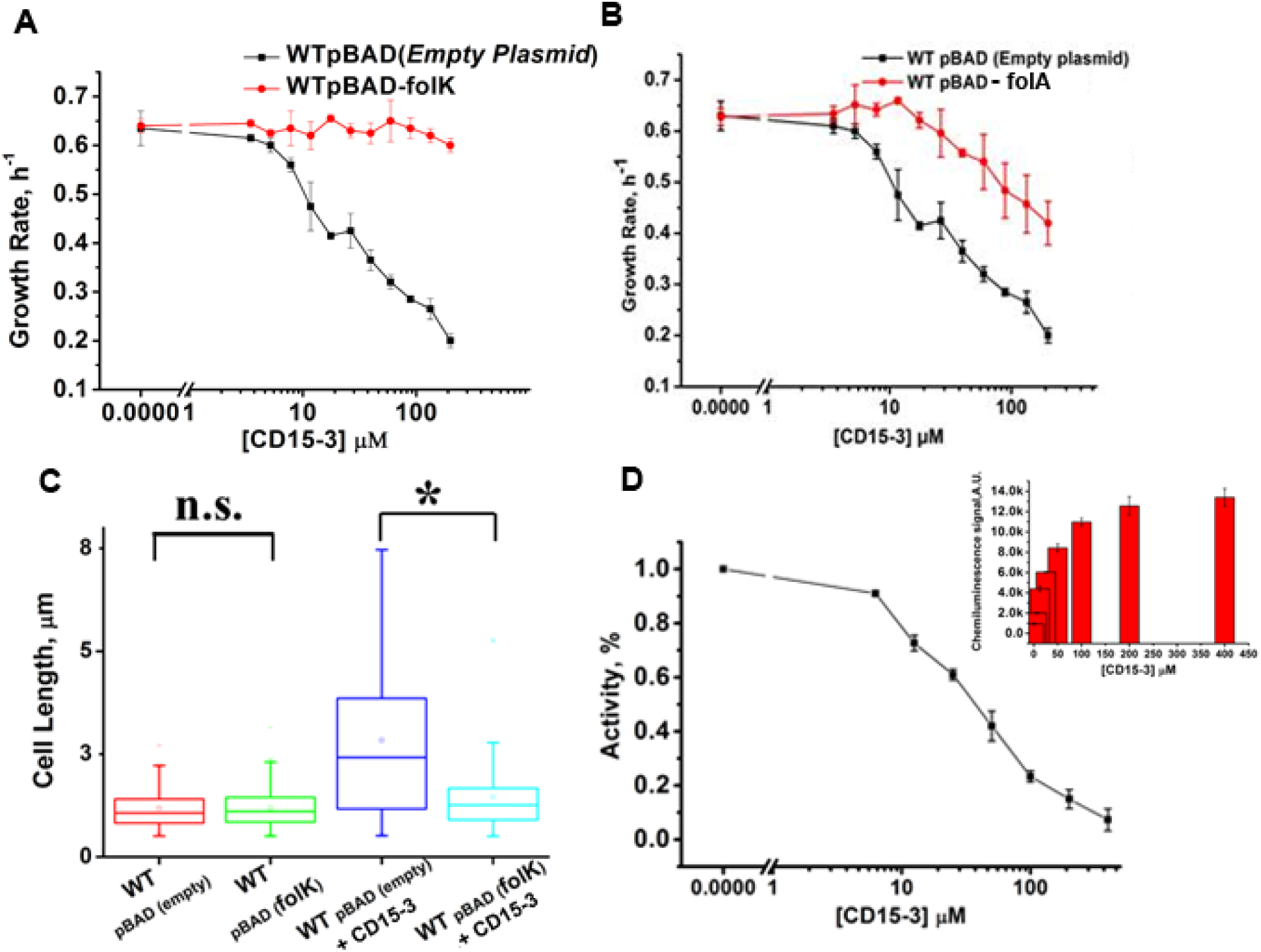
Overexpression of folK (encoding HPPK) rescued cells from CD15-3 induced growth inhibition. (A) Overexpression was induced with 0.1% arabinose and folK was expressed under pBAD promoter. folK rescued growth rate at all CD15-3 concentrations. (B) Overexpression of folA (encoding DHFR) was induced at 0.005% arabinose. folA overexpression had only partial effect in recovering growth rate in CD15-3 treated cells. (C) Distribution of cell lengths as derived from DIC imaging of WT and WT cells overexpressing folK under control and CD15-3 treatment conditions. Untreated WT and cells overexpressing folK had comparable cell lengths with median cell lengths of 1.06 µm. (n.s. indicates the distribution of cell lengths were not significantly different, Mann-Whitney test, p-value =0.952). Both WT and cells overexpressing folK were subjected to CD15-3 treatment at IC_50_ concentrations. WT treated cells had a filamentous morphology and the median cell length (2.41 µm) double of that of the untreated WT set. WT cells overexpressing folK after CD15-3 treatment had a median cell length of 1.252 µm which is slightly higher than that of untreated set. However, the cell size distribution of the cells overexpressing folK showed much less change after CD15-3 treatment compared to that observed for the WT (* indicates the distribution of cell lengths were significantly different, Mann-Whitney test, p-value <0.001). (D) Inhibition of HPPK (encoded by folK) by CD15-3 in an in vitro assay (see text and Methods for details). Residual activity of HPPK at various concentrations of CD15-3 Inset shows the bar plot for the chemiluminescence signals at various concentrations of CD15-3 in the HPPK assay buffer.

We further over-expressed folP (encoding DHPS) (Supplementary Figure 4A), folC (encoding DHFS) (Supplementary Figure 4B) and folD (encoding MTHFC) (Supplementary Figure 4C) to see any possible promiscuous rescue effect on growth rates. Only at lower concentrations of CD15-3 (<50µM) overexpression of these genes partially rescued growth inhibition. Overexpression of folD (Supplementary Figure 4C) showed slight improvement in growth rate at mid concentration of CD15-3 (around 70 µM). *E. coli* folD gene encodes for a bifunctional enzyme having both methylenetetrahydrofolate dehydrogenase and methenyltetrahydrofolate cyclohydrolase activities. The dehydrogenase and cyclohydrolase reversibly catalyze oxidation of N5,N10-methylenetetrahydrofolate to N5,N10-methenyltetrahydrofolate and the hydrolysis of N5,N10-methenyltetrahydrofolate to N10-formyltetrahydrofolate and play critical role in purine metabolism.

As an additional negative control, we overexpressed two more genes and found no recovery effect. The metF gene encodes for methylene THF-reductase (MTHFR). Overexpression of metF did not show any rescue in CD15-3 induced growth inhibition (Supplementary Figure 4D). Next, we overexpressed gene pabC which encodes for ADCL. pabC encoding aminodeoxychorismate lyase is involved in the biosynthesis of p-aminobenzoate (PABA) which is a precursor of tetrahydrofolate. ADCL converts 4-amino-4-deoxychorismate into 4-aminobenzoate (PABA) and pyruvate. Overexpression of pabC did not show any recovery of growth rates in CD15-3 treated cells (Supplementary Figure 4E).

Of all the candidate genes, only folK overexpression showed a clear rescue effect at all studied CD15-3 concentrations, with growth approaching WT level of around 0.6 hour^-1^ at all concentrations of CD15-3. This indicates strongly that CD15-3 interacts with cellular HPPK as its non-DHFR molecular target. The complete growth recovery observed with folK overexpression indicates that the growth-limiting folate perturbation originates in the folK mediated step, which in turn impacts rest of the folate pathway. In the Supporting Text (Supporting Information) we provide an explanation as to why folA overexpression leads only to partial rescue while overexpression of folK resulted in full recovery from CD15-3 induced inhibition. In short, the reason for the difference is in different expression levels of folA and folK in overexpression experiment. The Supporting Text presented quantitative analysis of the effects of inhibition of two proteins by a common inhibitor and the competing factors of inhibition of both enzymes and sequestration of the inhibitor by an overexpressed enzyme. Supplementary Figure 5 summarizes the effects of various metabolic supplements used and gene overexpression strategies deployed and show how CD15-3 has a folate-related mechanism of action.

Perturbation in the folate pathway leads to cellular filamentation and concomitant morphological changes (Ahmad et al., 1998; Bershtein et al., 2015; Bhattacharyya et al., 2021; Justice et al., 2008; Sangurdekar et al., 2011; Zaritsky et al., 2006). WT cells treated and untreated with CD15-3 were grown for 4 hours and subjected to DIC imaging. CD15-3 treated cells (Supplementary Figure 6B) showed a considerable extent of cellular filamentation as compared to untreated WT (Supplementary Figure 6A) cells grown for the same length of time. A similar experiment was also done with WT cells overexpressing folK under pBAD promoter with 0.1% arabinose induction. WT cell sets overexpressing folK were grown for 4 hours under control and CD15-3 treatment conditions. WT cells overexpressing folK did not show any visible change in cellular shape and size (Supplementary Figure 6D) compared to the untreated control (Supplementary Figure 6C). Upon comparison of the median cell lengths (Figure 6C), a slightly higher median cell length was observed in the folK overexpressing cells with CD15-3 treatment (median cell length = 1.252µm), as compared to untreated cells under control conditions (median cell length = 1.06µm). This slightly higher median cell length could be attributed to the fact that remaining CD15-3 unsequestered by HPPK also targets cellular DHFR. Overexpression of folK although mostly reverses the effects of CD15-3 on cell shape; The overall pronounced rescue in cell length upon overexpressing folK further supports the conclusion that HPPK is the non-DHFR cellular target of CD15-3.

### In vitro assay confirms CD15-3 is an inhibitor of HPPK encoded by folK

Next, we aimed to verify in an *in vitro* assay that CD15-3 indeed inhibits HPPK in a concentration dependent manner. To that end we performed a KinaseGlo^TM^ assay to test for HPPK activity and its probable inhibition in presence of CD15-3. We derived HPPK by 1000 fold dilution of the cell lysates overexpressing HPPK under pBAD promoter at 0.1% arabinose induction. KinaseGlo^TM^ assay is based on chemiluminescence (Tanega et al., 2009). With higher ATP concentration in the assay buffer, luciferase leads to the conversion of beetle luciferin to oxy-luciferin with the emission of light. The HPPK mediated reaction utilizes ATP leading to ATP depletion and hence drop in the chemiluminescence signal. Any potential inhibition of HPPK would retain original concentration of ATP keeping the chemiluminescence signal intact or like the control (with no HPPK activity). We observed a marked drop in the absolute chemiluminescence signal intensity in the HPPK reaction set with no CD15-3 in the assay buffer (Figure 6D inset). Interestingly presence of CD15-3 led to enhanced absolute chemiluminescence signal intensity (Figure 6D inset) suggesting that CD15-3 does inhibit HPPK. The inhibitory effect appeared to be progressively higher upon increase of the CD15-3 concentration in the reaction-assay-buffer. For control we performed similar experiments with 400 µM of CD15-3 with one ATP dependent protein Adk and one ATP independent protein BSA to validate that the drop in signal we observe in presence of CD15-3 in the HPPK reaction-assay set is due to the specific inhibitory interaction between CD15-3 and HPPK (Supplementary Figure 7). We did not observe any drop in luminescence with BSA or ADK reaction set in the presence of CD15-3.

In our assay protocol, HPPK reaction was initiated with the introduction of the substrate, 6-Hydroxymethyl-7,8-dihydropterin in the assay buffer (Supplementary Figure 7). We used the absolute chemiluminescence intensity values to calculate % activity using the following relation:

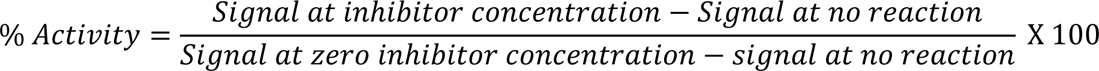

where signal at inhibitor concentration is the chemiluminescence signal at any given concentration of CD15-3, signal at no reaction is the optical signal obtained when the substrate is not added, and the reaction is not initiated.

Using this relation and plotting the absolute signal intensity values observed across the CD15-3 concentration gradient we found that HPPK retains 50% activity at 36.46 µM (IC_50_) (Figure 6D) showing that CD15-3 is indeed an inhibitor of folK.

## Discussion

Intracellular drug target identification is a hard problem. Often candidate drugs interact with unintended proteins inside cells and the resultant phenotypic effect emerges from the off-target protein(s). This applies to drugs spanning from antibiotics (Silver, 2007) to anti-cancer formulations (Lin et al., 2019). A systematic understanding of cellular targeting is critical in drug discovery programs as it provides mechanistic insights into intracellular drug action. This understanding in particular stands critical in the context of drug resistance, as drug-resistant cells can mount plethora of strategies to evade the drug action. For example, the bacterial “resistome” is a tight assembly of multi-layered highly orchestrated mechanisms (Olivares Pacheco et al., 2013). In the current context of widespread antibiotic resistance, including the emergence of “multi-drug resistant” ESKAPE variants (De Oliveira et al., 2020), a mechanistic understanding of intracellular antibiotic targeting and what leads to bacterial death stands as an immensely pertinent problem.

Previously we reported CD15-3 as a potential antibiotic that significantly constrains bacterial evolvability by blocking the evolutionary escape routes which *E. coli* traverses under Trimethoprim selection (Zhang et al., 2021). We hypothesized that CD15-3 is an interesting lead towards development of the evolution drugs to overcome antibiotic resistance. However, our in-cell experiments showed that CD15-3 has an additional unidentified non-DHFR target; thus, blocking more than one cellular target makes evolution of antibiotic resistant *E. coli* phenotypes more difficult. It was important to determine the alternative target of CD15-3 in *E. coli* cells to better understand the mechanism of action of this prototypic “evolution-drug” inside bacterial cells. Using CD15-3 as a test case, in the present study we developed an integrated multi scale systems guided framework utilizing global metabolomics interpreted through machine learning and metabolic modelling, gene overexpression assays, and growth recovery studies eventually analysing data in the context of the metabolic network to unravel the unknown intracellular antibiotic target.

Antimetabolite classes of antibiotics such as antifolates (for example Trimethoprim) target proteins at critical points in the bacterial metabolic scheme. Hence an investigation of the metabolic architecture provides essential clues tracing the potential points of metabolic perturbations under conditions of antibiotic action. Critical analysis of such points of metabolic perturbation and its comparison with untreated control sets provides mechanistic insights of the drug action inside cell. Recent advances in untargeted metabolomics have provided valuable insights into the global metabolome and helped to quantitatively identify the metabolic cascades impacted by perturbations (Vincent et al., 2016; Wu et al., 2015; Zampieri et al., 2017). Also, machine learning methods have become increasingly popular for statistical analysis of the metabolomics data due to the inherent non-linear metabolomic data representation and the ability to process large and heterogeneous data rapidly (Bagherian et al., 2021; Liebal et al., 2020). While model interpretation has been a historical challenge in deploying machine learning for biological data analysis, interpretable ‘white box’ machine learning methods have come into focus as a viable area of development to empower drug discovery workflows (Yang et al., 2019).

Employing machine learning on the comparative metabolomics data and training a multi-class K-nearest neighbour model, we found that CD15-3 induced metabolomic perturbation has a typical antifolate signature, suggesting that the unknown target is located somewhere in the folate pathway. Focusing on folate pathway perturbation and performing metabolic supplementation experiments we observed that supplementation with a subset of metabolites lead to growth recovery. Our analysis in the context of the metabolic network utilizing constraint-based metabolic modelling confirmed that inhibition of folate metabolism was consistent with patterns of growth rescue. We utilized protein structural analysis to suggest targets upstream of the DHFR catalysed step within the folate pathway and performed gene over-expression studies for these target-genes to determine which candidate targets rescue growth inhibition by CD15-3. Among all the candidate genes folK, which codes for HPPK, showed complete rescue of growth rate under CD15-3 treatment conditions. Unlike DHFR overexpression which only partially rescued CD15-3 induced growth inhibition (Yang et al., 2019), HPPK showed clear recovery at all concentrations of CD15-3. In the Appendix we discuss a plausible explanation as to why we see a full rescue from CD15-3 induced growth inhibition with HPPK overexpression compared to partial rescue observed with DHFR overexpression.

This work demonstrates a promising path towards white box machine learning workflows through coupling standard machine learning modelling with mechanistic metabolic modelling and protein structural analysis. As the release of AlphaFold (Varadi et al., 2022) is making protein structure prediction an increasingly accessible task, this approach utilizing protein structural comparison could provide a new direction to rapidly demystify drug target identification (DTI).

With HPPK as the non-DHFR target, CD15-3 can be considered as the multivalent drug which can simultaneously block and inhibit two molecular targets (DHFR and HPPK) inside the bacterial cell. Interestingly, being an essential protein, bacterial HPPK has been an attractive target for designing antibiotics (Shaw et al., 2014). Thus CD15-3 in principle can serve as a lead to a “monotherapy-analog” of “combination therapy” which blocks the emergence of antibiotic resistant phenotypes by interacting with two targets making it difficult for bacteria to escape antibiotic stress.

We showed how integrating constraint based metabolic modelling with machine learning in analysing large scale metabolomics data can help capturing metabolic perturbation signatures and narrow down the search options for identifying intracellular drug target. We acknowledge that one limitation of the current proposed framework is this that it works with antimetabolite classes of antibiotics and has not been tested with antibiotics in general. Future work would aim at development of similar workflow for other classes of antibiotics where perturbations remain highly delocalized and impacts non-metabolic cellular functions. Further, the problem of off-target activity is also highly relevant for drugs beyond antibiotics. Many anti-cancer drugs have been reported to have off-target toxicities (Lin et al., 2019). We believe our proposed framework would be relevant in addressing these off-target identification problems. Future works would involve using this multi-scale framework for off-target identifications in other cellular models and complementing it with relevant functional assays.

## Material and Methods

### Antibacterial Growth Measurements and IC_50_ values

Bacterial cultures in M9 minimal medium were grown overnight at 37 °C and were then normalized to an OD of 0.1 using fresh medium. A new normalization to an OD=0.1 was conducted after additional secondary growth for ∼4 hours. Thereafter the M9 medium and six different concentrations of the CD15-3 in the 96-well plates (1/5 dilution) were incubated. The incubation of the plates was performed at 37 °C and the orbital shacking and absorbance measurements at 600 nm were taken every 30 min during 15 h. Growth rate was calculated using logistic fitting on matlab.

### Metabolomics analysis

Untargeted global metabolomics was performed to understand the global metabolome of the WT treated and control sets under different experimental contexts. In all the experiments cells were grown in M9 minimal media supplemented with 0.8g/L glucose in a 250 mL flask and temperature of 37^0^C was maintained. Cells were pelleted with a brief 2 minutes pre-centrifugation incubation step on dry ice. After pelleting the cell pellets were mixed with 80% pre-chilled methanol Samples were thereafter vortexed and incubated in dry ice for 10 min followed by centrifugation at 4 °C for 10 min at maximum speed. The supernatant was collected, and the pellet was repeatedly processed by resorting to the above-mentioned procedures. Samples were stored at −80°C until analyzed by mass spectrometry. In all our experiments at least three independent biological replicates were analyzed. LC-MS analysis in the positive and negative mode was performed as previously described (Bhattacharyya et al., 2016). A list of 48 experimentally measured retention times was used for initial calibration of the retention time predictions. We performed data analysis for untargeted metabolomics using the software the packages MzMatch (Scheltema et al., 2011) and IDEOM (Creek et al., 2012). In untargeted analysis for peak assignment IDEOM include both positive and negative peak M/Z values and predicted retention times calculated based on chemical descriptors. We followed the same method of analysis as we had in one of our earlier studies (Rodrigues and Shakhnovich, 2019b).

### Recovery Experiments

WT BW25113 cells were grown both in absence and presence of prospective compound CD15-3 in M9 media supplemented with 0.8gL^-1^ glucose. Both the naïve (which were not exposed to CD15-3 treatment) and pre-exposed cells (cells treated with CD15-3) were subjected to growth in normal M9 media and growth profiles were analyzed. The entire pre-exponential stage (lag phase) was grouped into equal time frames and both naïve and pre-exposed cells were harvested for metabolomic studies. Same process was also executed for harvesting both naïve and pre-exposed cells at late log phase.

### Overexpression Experiments

WT cells were transformed with blank vector plasmids (without inserts) as well as plasmids overexpressing genes viz. thyA, glyA, metF,purH, purC,folD,purD,adk and folK. The genes were under pBAD-promoter and the overexpression of the genes was induced using externally supplemented arabinose (0.1%).

### Machine learning analysis of metabolomics data

Metabolomics data for ten antibiotics was taken from a published study (Zampieri et al., 2017). The data was first filtered based on the criteria of having annotated, high confidence identities (annotation score > 50) and overlap (shared KEGG identifier) with the data generated in this study for CD15-3. Data was then averaged for each compound. Uniform Manifold Approximation and Projection (UMAP) was utilized to visualize the high-dimensional data in two dimensions, to inspect clustering behavior of the samples. A multi-class logistic regression model (Pedregosa et al., 2011) was trained to identify drug mechanism of action from metabolomics data. Five mechanisms of action were utilized from the original study: antifolates, cell wall synthesis, polymerase inhibition, translation inhibition, and oxidative stress. Mechanism labels were utilized as target values in a supervised learning approach. Zero time point data was excluded. UMAP and the LR algorithm were implemented using the scikit-learn Python package (Pedregosa et al., 2011). UMAP was implemented with 2 components, fixed random state of 42, and 14 neighbors, based on the number of samples for high and low concentration for a single drug. The LR algorithm was implemented with default parameters, and performance was evaluated with leave-one-out cross-validation.

### Metabolic modeling analysis of metabolomics data

To analyze patterns in growth rescue data from metabolic supplementation, we utilized a flux balance analysis workflow (Orth et al., 2010). We utilized MATLAB, the COBRA toolbox version 3.0 (Heirendt et al., 2019) and the latest metabolic network reconstruction of E. coli, termed iML1515 (Monk et al., 2017). Default options were used for the optimizeCbModel function.

To enforce folate limitation, growth was first optimized under glucose growth at an arbitrary but realistic uptake rate of 10 mmol/gdW/hr. Then, folate dependent reactions nearby to each metabolite supplement were identified. Reactions that consumed folate were limited to 90% of their optimal flux, and it was confirmed that growth rate was correspondingly limited. Then, metabolites were supplemented in silico at a rate of 0.1 mmol/gDW/hr. The calculated growth rate following supplementation was compared to the calculated growth rate before supplementation to determine the growth benefit of the metabolite under folate limitation.

Gene expression data was used to close flux through reactions that were not expressed under wild type conditions in a targeted manner around metabolites that were supplemented. Closing flux through all reactions that were not measured to not be expressed in the proteomics data was not possible due to lack of growth in the resulting simulations.

The resulting scores were combined for different pathway specific inhibitions for each supplement, as detailed in the code available in the Supplementary Materials.

For non-folate pathway comparisons, reactions were constrained to 0 flux, and the benefit of each metabolite supplement for each reaction inhibition was calculated.

### Protein structural comparison

For the global structural analysis, general protein structural properties were taken from the *E. coli* GEM-PRO for all available metabolic enzymes. These properties were calculated previously as part of the GEM-PRO development. These structural data was standard normalized and then correlated with Pearson correlation to examine global structural similarity.

In the targeted folate pathway structural analysis, proteins were selected that were nearby in metabolic maps in the pathway of interest, as well as some unrelated metabolic proteins as a control comparison. Isomif (Chartier et al., 2016) was used to pull different binding clefts from each of the selected proteins and the 5 largest of each protein binding clefts were kept. The similarity (Tanimoto coefficient) between the clefts of the folA protein structure (PDB ID: 1DRA) and each of the selected clefts of the proteins using isomif which was then plotted to view which binding clefts were most similar. FATCAT (Li et al., 2020b) was used to compare the overall structural similarity between all of the selected proteins and the results were plotted using a clustermap to look for similar proteins. Visualizations of the proteins were created using nglview.

### Statistical and Python plots

Statistical analyses of the data and their representation was carried out using Origin pro 8.1 package. Metabolomics data were processed using R based MS converting operation and IDEOM tool. For calibrating M/Z (mass to charge) values and retention times of the standard metabolites, XCalibur package was used. For the quantitative depictions of the metabolomics data statistically validated outputs were plotted using Python libraries of matplotlib and seaborn.

### HPPK activity assay

HPPK activity assay was conducted with the help of KinaseGlo^TM^ assay kit. In this assay, firefly luciferase utilizes the ATP remaining after HPPK catalysis to produce a luminescence signal that is directly proportional to ATP concentration; from this, the HPPK activity can be derived. The enzyme activity calculation and selection of optimum concentration was done following previously published methods (Chhabra et al., 2012). For kinetic measurements, an optimized HPPK concentration of 7 ng/50 µL assay volume was determined, which allowed for monitoring the first 10% of reactions turnover in a reasonable assay time period (20 min).

Measurements were performed in 96-well plates using assay buffer (100 mMTris-HCl/10 mM MgCl_2_, pH 8.5, 0.01% (w/v) BSA, 0.01% (v/v) Tween 20 and 10 mM β-mercaptoethanol). Typically, 5 µl of test compound (dissolved in 50% DMSO) and 20 µl of enzyme were added to each well followed by 25 µl of assay buffer giving 0.3 µM pterin and 0.2 µM ATP in a total reaction volume 50 µl. After a 20 minute incubation at room temperature, the enzymatic reaction was stopped with 50 µl of KinaseGlo™ reagent. Luminescence was recorded after a further 10 min using the plate reader (Tecan Infinite M200 Pro).

### Differential interference contrast (DIC)

WT cells were grown in M9 media supplemented with 0.8gL^-1^ glucose and casamino acids (mixtures of all amino acids except tryptophan) in absence and presence of CD15-3 at 42^0^C for incubation and 300 rpm constant shaking. Drop in DHFR activity has been associated with cellular filamentation and similar phenotype is observed under TMP treatment(Bhattacharyya et al., 2021). Since CD15-3 targets intracellular DHFR and soluble fraction of cellular DHFR is lower at 42 degrees C we chose this temperature for our imaging studies (Bershtein et al., 2012).

Aliquots were taken from the growing culture and they were drop casted on agar bed/blocks. These blocks were taken further processed for differential inference contrast (DIC) imaging using Zeis Discovery imaging workstation. Multiple fields were observed and scanned for a single condition type and a minimum of three replicates were used for imaging studies. Similar methods for imaging were used for WT cell types overexpressing folK under conditions of absence and presence of CD15-3 compound. Intellesis Module was used for analyzing DIC images. On average, around 500 cells were analyzed for computing cell length. *E. coli* cell lengths in our imaging studies were not normally distributed. Nonparametric Mann-Whitney test was therefore used to determine if the cell length distributions were significantly different upon CD15-3 treatment.

## Appendix: The case when 2 enzymes are inhibited by a common inhibitor

Here we consider the situation when two enzymes are inhibited by common inhibitor. We use the following notations:

*x*_0_ total concentration of the inhibitor (i.e. free + bound)

*x_free_* concentration of free inhibitor in solution

*E*_1_^0^ total concentration of enzyme 1 in the cell

*E*_1_^*free*^ concentration of free (unbound) enzyme 1 in the cell

[*E*_1_*x*] concentration of bound enzyme 1-inhibitor complex in the cell

*E*_2_^0^ total concentration of enzyme 2 in the cell

*E ^free^* concentration of free (unbound) enzyme 2 in the cell

[*E*_2_ *x*] concentration of bound enzyme 2-inhibitor comnplex in the cell

K_*d*1_, K _*d*2_ binding affinities of enzyme 1 and 2 respectively to the inhibitor

Using Law of Mass Action and conservation of mass for all three components (enzymes 1 and 2 and the inhibitor) we get for both enzymes:

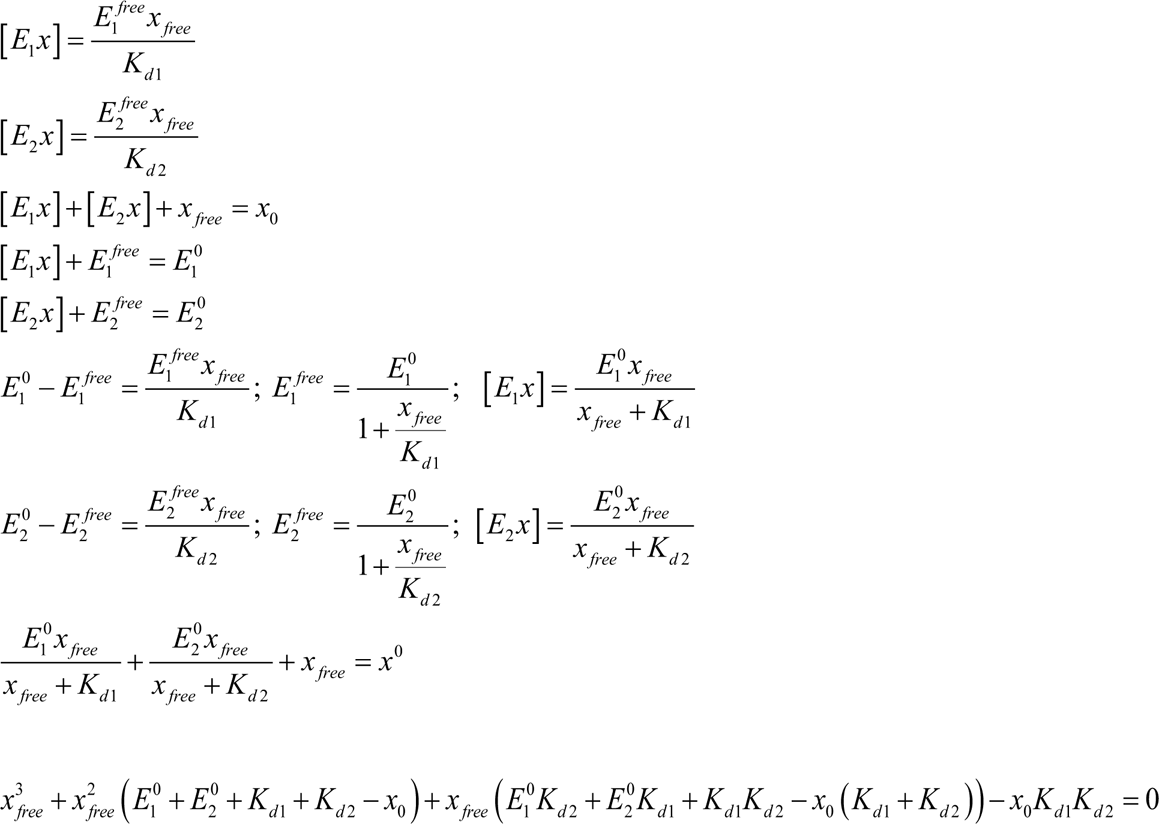

Now consider the overexpression case 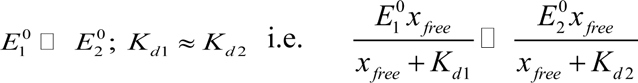

This inequality means that most of the inhibitor is bound to enzyme 1 (naturally as it is overexpressed). In this case the cubic equation for free concentration becomes quadratic because we neglect in the first approximation the inhibitor bound to enzyme 2

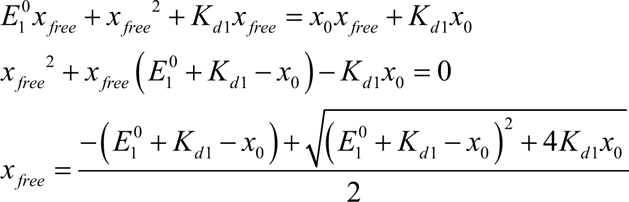

assume high overexpression

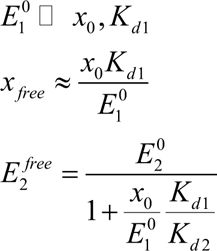

Thus, the effect of overexpression of one enzyme affects the way how another enzyme is inhibited by the same inhibitor in the positive way: the more one enzyme is overexpressed the less other enzyme is inhibited because the inhibitor is sequestered into a binding complex with the overexpressed enzyme. However, with respect to increased expression of the inhibitor we can see 2 regimes:

when overexpression is high 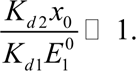

In this case 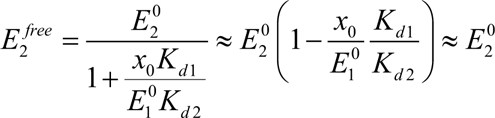

i.e. fitness does not depend on the inhibitor concentration because the overexpressed enzyme sequesters most of the inhibitor and the other target enzyme can function uninhibited.

In the opposite case of high concentration of the inhibitor relatively to the overexpressed enzyme or when binding affinities differ strongly between two enzymes, residual inhibitor can inhibit second (not overexpressed) enzyme.

This scenario corresponds to the case 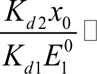 i.e., when there is plenty of inhibitor - more than overexpressed enzyme can bind - the approximation is different, and the result is

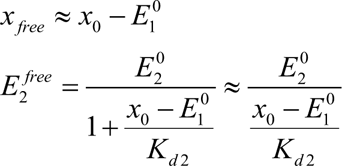

We conclude that there can be an asymmetry between the effect of overexpression of HPPK and DHFR. If HPPK is highly overexpressed such that it sequesters most of the inhibitor, then fitness – concentration of the inhibitor curve will be flat. DHFR overexpression from pBAD was restricted to 0.005% arabinose induction to avoid overexpression induced toxicity (Bhattacharyya et al., 2016) which would complicate interpretation of the rescue experiments. If DHFR is less expressed from the plasmid, i.e. at lower concentration of arabinose as in our experiment (0.005% for folA vs 0.1% for HPPK) then upon addition of inhibitor the inhibitor-growth will be flat at lower concentration of the inhibitor followed by loss of fitness at higher concentration of the inhibitor due to inhibition of HPPK by remaining inhibitor. Inhibition of growth occurs in this case when concentration of the inhibitor becomes greater or equal to concentration of the overexpressed protein. E.g., 1000-fold overexpression of DHFR equals to about 100 micromoles concentration in the cell – roughly similar to concentration of CD15-3 in the experiments.

## Supplementary figures

**Supplementary Figure 1:**
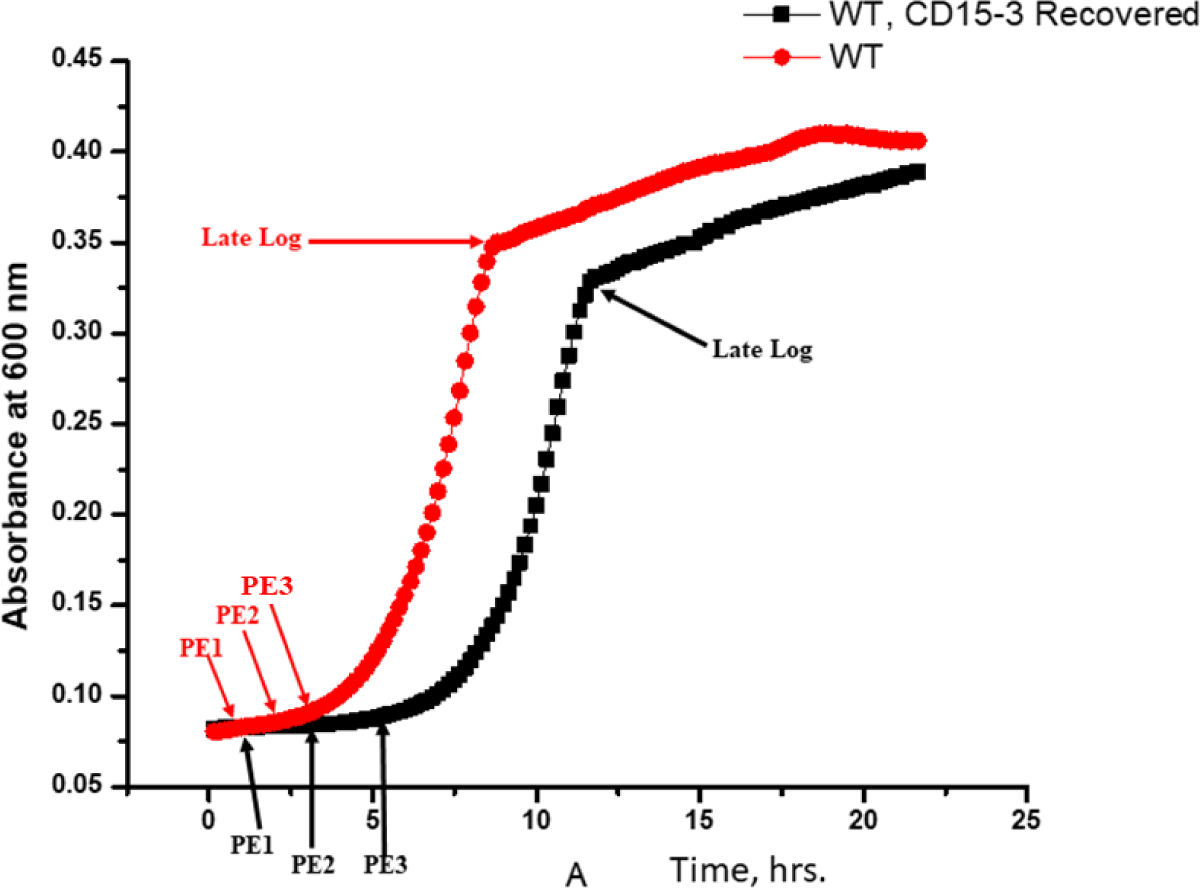
Growth curves for WT (naïve in red) and WT pre-exposed cells (CD15-3 recovered in black). PE1, PE2 and PE3 are respectively pre-exponential stages 1, 2 and 3 of growth curve. Stages (PE1, PE2, PE3 and Late Log) marked in red are for naïve WT cells and those marked in black refer to pre-exposed WT. For the recovery assay experiments, naïve and CD15-3 pre-exposed cells were harvested at their respective pre-exponential and late log phases of growth.

**Supplementary Figure 2:**
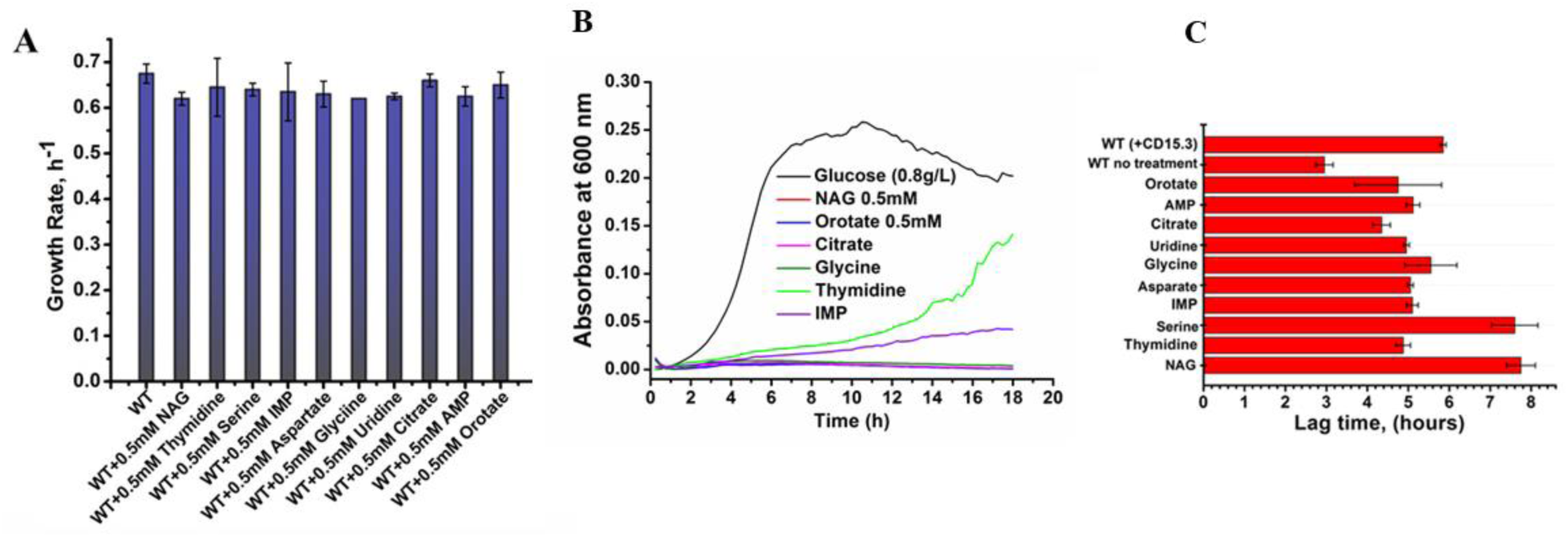
Metabolites selected for external supplementation experiments did not show any toxic effect on the growth profiles of the cells. (A) Bar plot showing growth rates of WT cells grown in the presence of metabolites as external supplements in M9 media. (B) Growth curve shows that these external metabolites did not show any potential to function as an alternate C-source in the growth media. (C) Bar plot showing variation in lag-time during metabolite supplementation experiments. NAG and serine supplementation leads to a prolonged lag time compared to other metabolites.

**Supplementary Figure 3:**
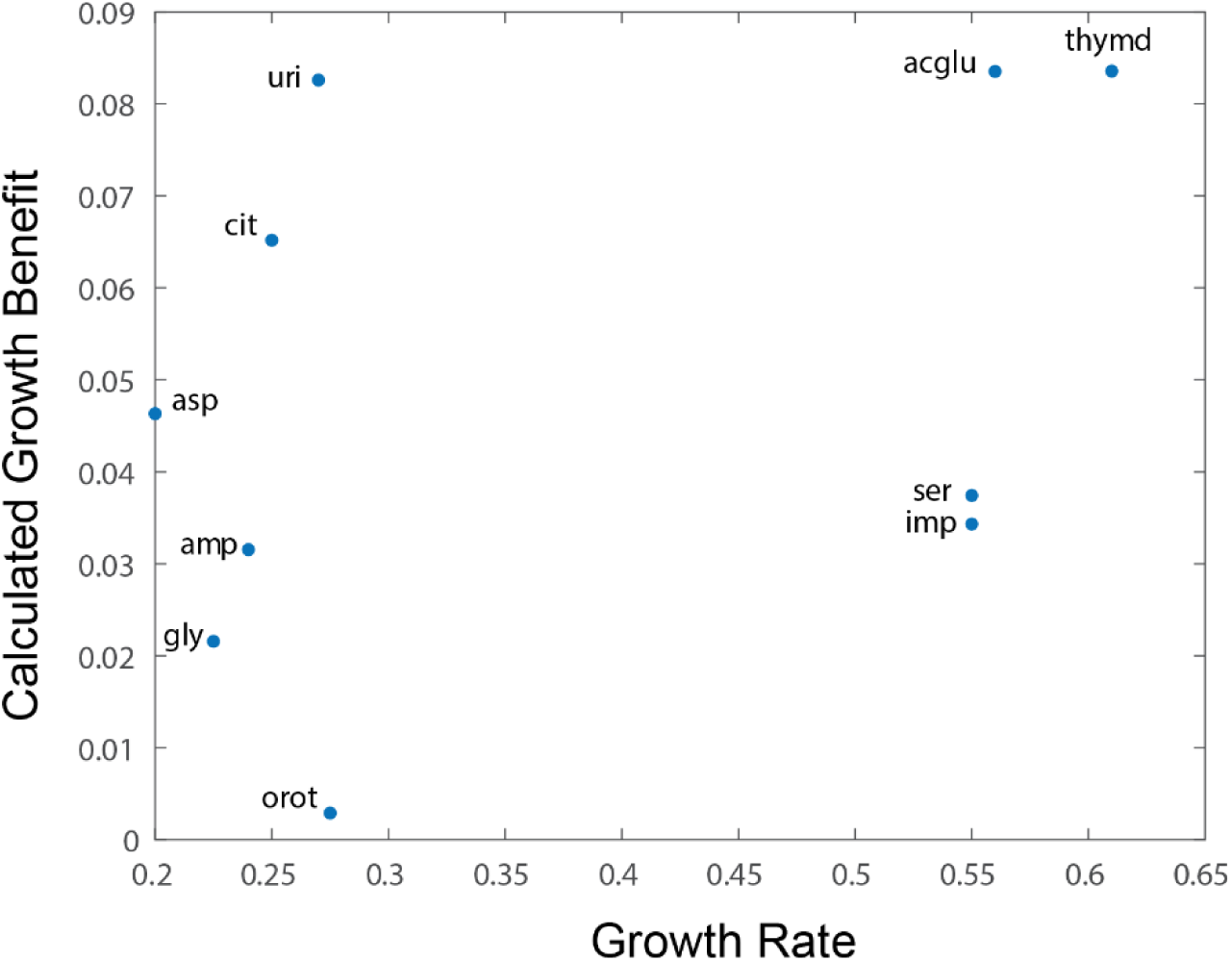
Correlation of metabolic supplementation induced growth rescue under CD15-3 treatment with model calculated growth benefit. Flux balance analysis was used to calculate a maximum growth flux solution under glucose growth, and solution shadow prices were extracted for each metabolite. Shadow prices represent the benefit of each metabolite to the FBA objective, which is growth. An overall correlation is observed, including for metabolites not listed in the main text, namely aspartate and n-acetyl glutamate, demonstrating a rationale for the ability of these metabolites to rescue growth

**Supplementary Figure 4:**
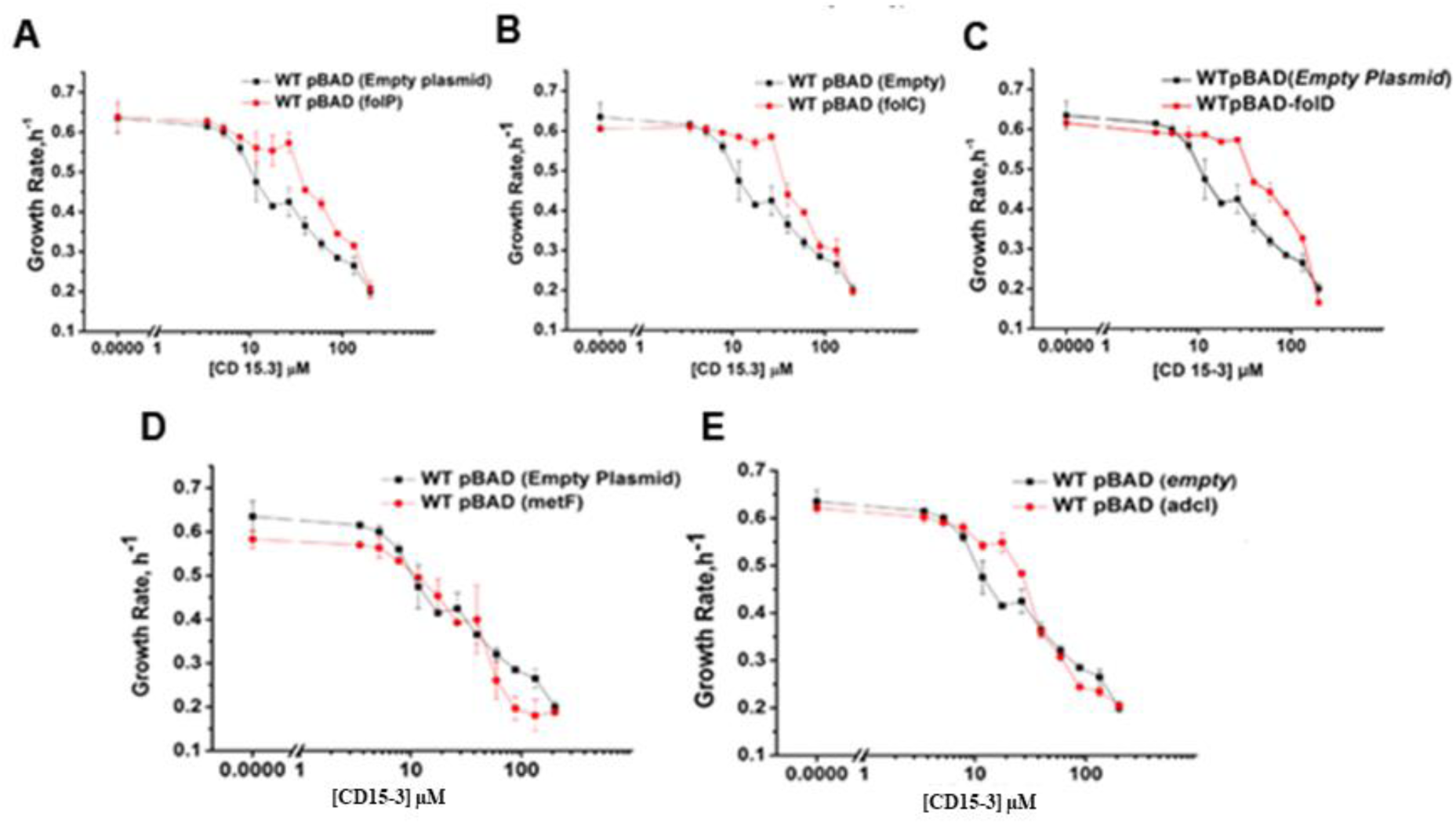
Overexpression of (A) folP (encoding DHPS) (B) folC (encoding DHFS) and (C) fol D (encoding MTHFC) showed slight improvement in the growth rates of the CD15-3 treated cells only at lower concentrations (<50 µM) of CD15-3. folP, folC and folD genes were overexpressed at 0.1% arabinose induction under pBAD promoter. (D) Overexpression of metF (encoding MTHFR) under pBAD promoter with 0.1% arabinose induction did not lead to recovery from CD15-3 induced growth inhibition. (E) Overexpression of ADCL (encoded by gene pabC) under pBAD promoter with 0.1% arabinose induction was found to have no recovery effect on CD15-3 treated cells

**Supplementary Figure 5:**
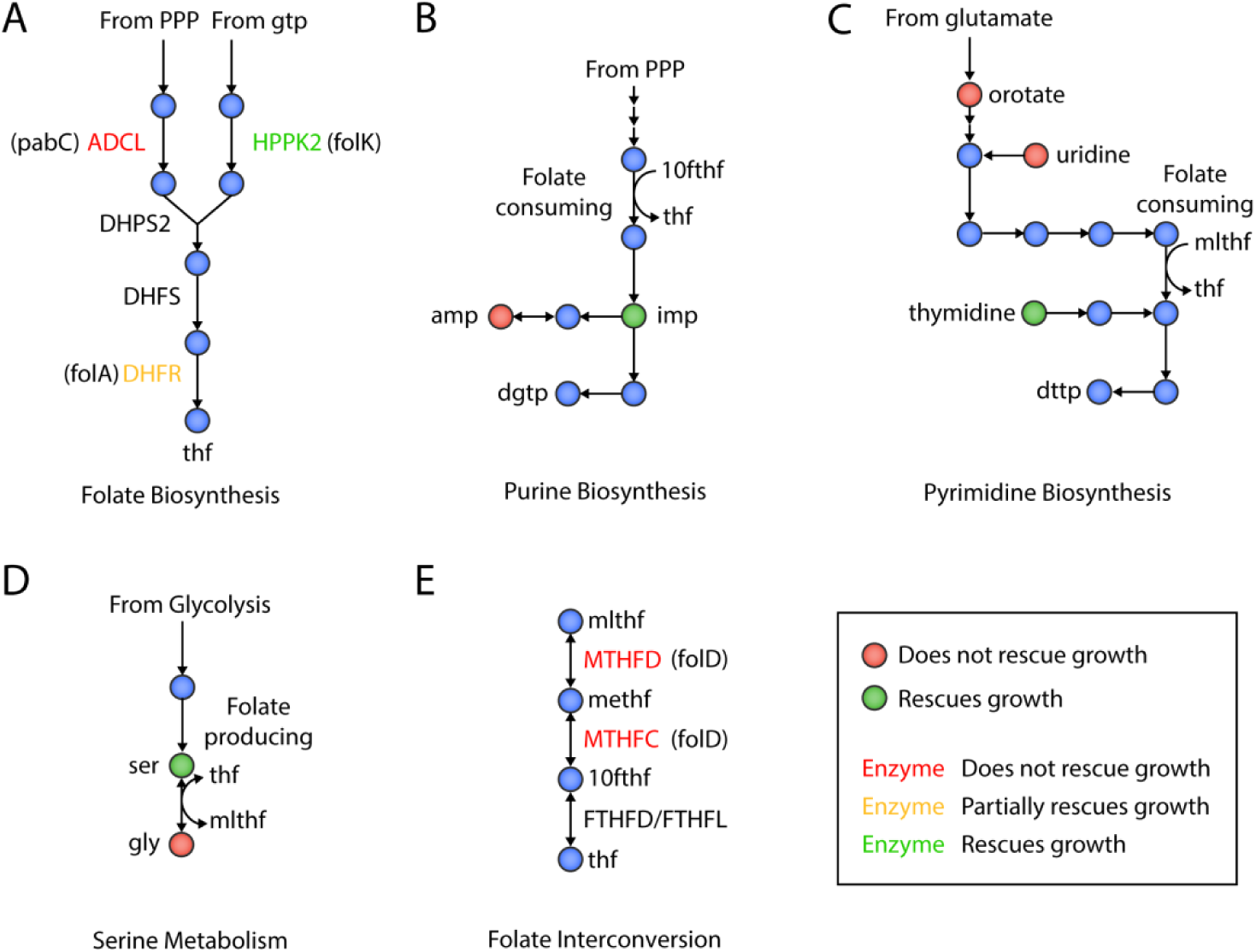
Summary of selected supplement and overexpression evidence for a folate-related mechanism of action of CD15-3. A) De novo folate biosynthesis. Three enzymes were overexpressed. Overexpression of the intended targeted, folA, partially restored growth. Overexpression of pabC (adcl) did not restore growth. B) Purine biosynthesis. IMP and AMP metabolic supplementation was done. IMP supplementation rescued growth while AMP had no effect in rescuing CD15-3 induced growth inhibition. C) Pyrimidine biosynthesis. Thymidine supplementation led to growth rescue from CD15-3 induced inhibition and Orotate and uridine had no effect in recovery. D) Serine biosynthesis. Metabolic supplementation of serine resulted in growth rescue. E) Folate interconversion.

**Supplementary Figure 6:**
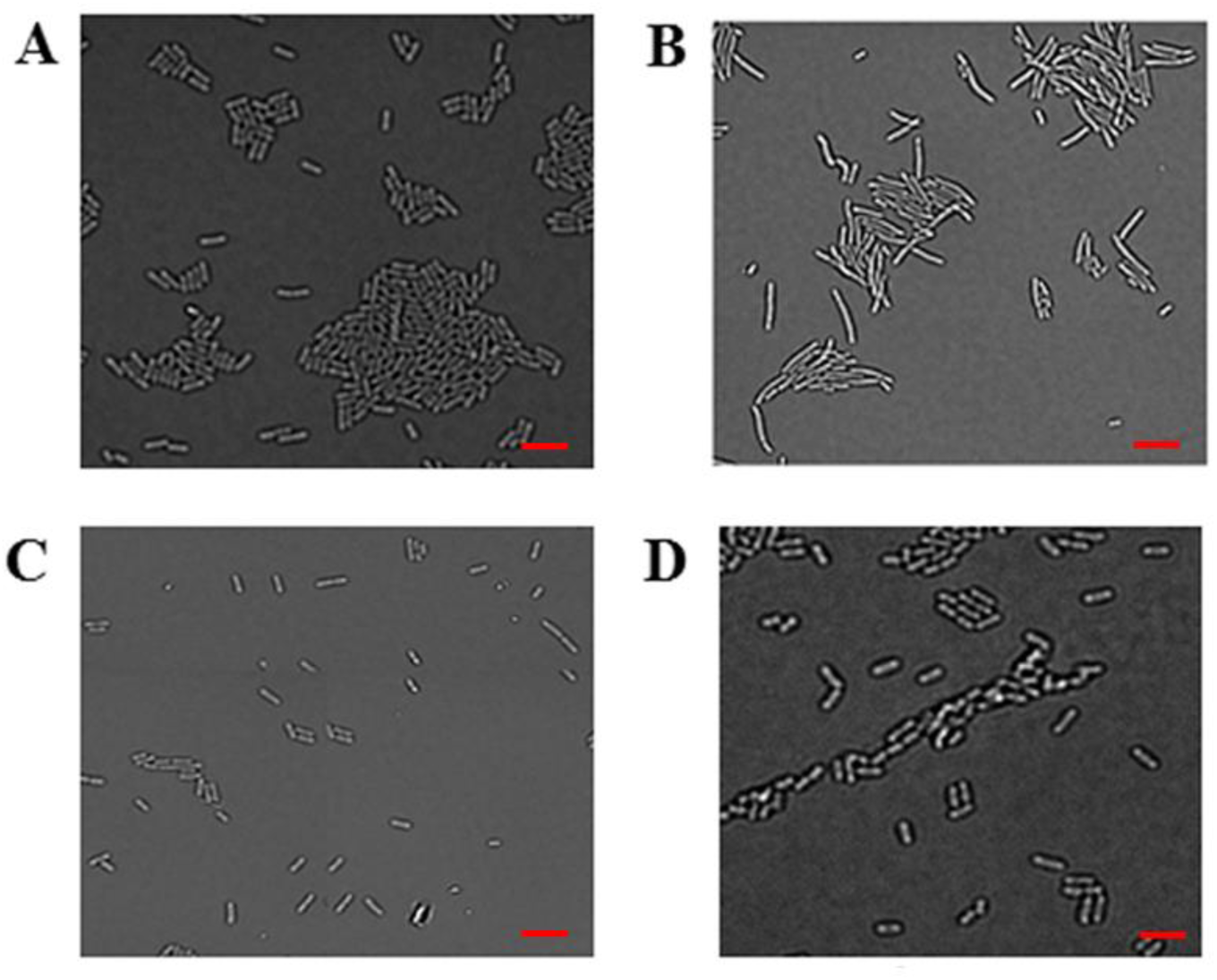
Overexpression of folK rescues cells from CD15-3 induced morphological changes. DIC image of WT cells under (A) control (no CD15-3 treatment) and (B) treated (CD15-3 treatment) conditions. CD15-3 treated cells shows visible signs of cellular filamentation. (C) DIC image of WT E. coli cells overexpressing folK under control (no CD15-3 treatment) and (D) CD15-3 treated condition. Unlike WT cells (B) showing cellular filamentation in presence of CD15-3, folK overexpressing cells upon CD15-3 treatment did not manifest visible signs of filamentation. Scale bar corresponds to a cell length of 2 µm.

**Supplementary Figure 7:**
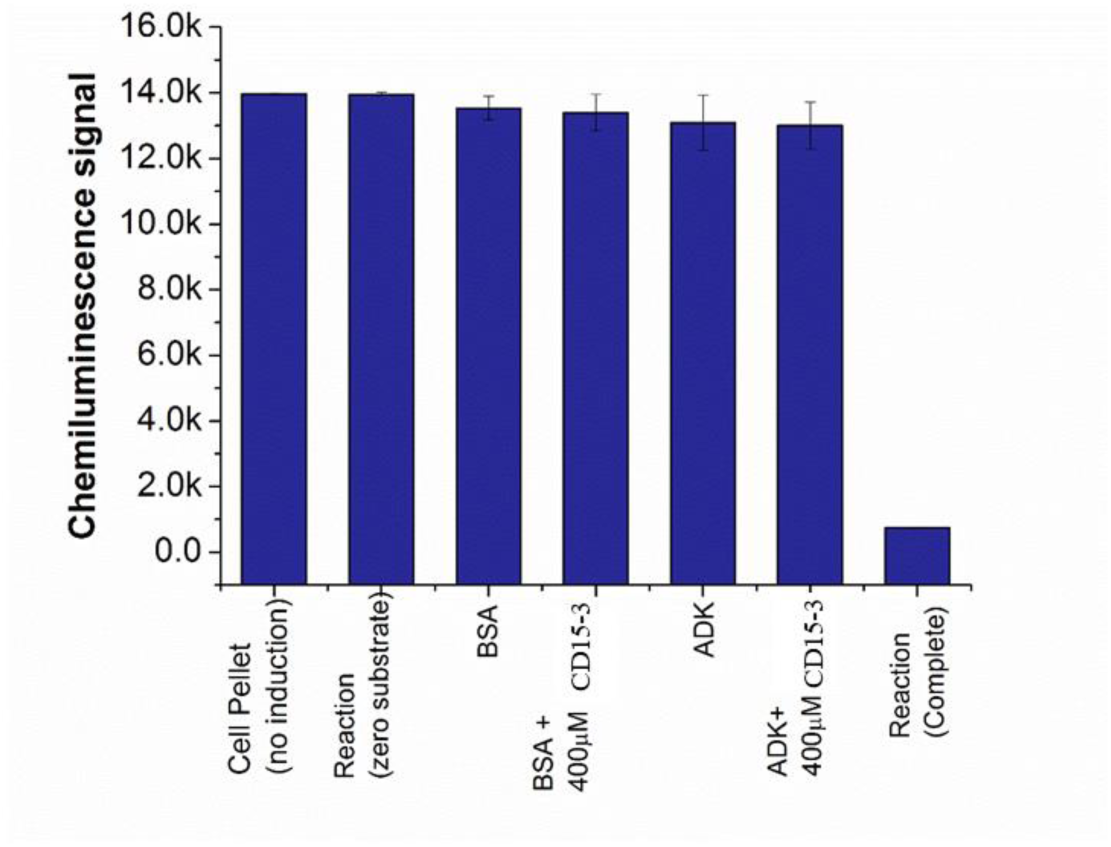
Bar plot showing the range of chemiluminescence signal at zero substrate (no reaction) condition and the end of reaction. Absolute chemiluminescence signal intensity as observed under different control conditions. 1000 fold diluted cell pellets (same as is used for folK overexpressing cells) obtained from cells not induced with 0.1% arabinose did not lead to any drop in the chemiluminescence signal. Similar readout was also found with BSA (a non-ATP using protein) and Adk (an ATP dependent protein) in presence of CD15-3 in the assay buffer.

## Acknowledgement

This work is supported by NIH R35GM139571 (to E.S.). We are grateful to Amir Bitran for helping in the omics data analysis, Sanchari Bhattacharyya for providing some of the plasmid constructs and Bharat V. Adkar for his help in the *in vitro* enzyme assay optimization.

